# Nutritional sex-specificity on bacterial metabolites during mosquito development leads to adult sex-ratio distortion

**DOI:** 10.1101/2023.10.18.562973

**Authors:** Ottavia Romoli, Javier Serrato-Salas, Chloé Gapp, Yanouk Epelboin, Pol Figueras Ivern, Frédéric Barras, Mathilde Gendrin

**Author notes:** These authors contributed equally and are listed by alphabetical order (Ottavia Romoli, Javier Serrato-Salas).

## Abstract

Mosquitoes rely on their microbiota for B vitamin synthesis. We previously found that *Aedes aegypti* third-instar larvae cleared of their microbiota were impaired in their development, notably due to a lack of folic acid (vitamin B9). In this study, we found that diet supplementation using a cocktail of seven B vitamins did not improve mosquito developmental success, but rather had a significant impact on the sex-ratio of the resulting adults, with an enrichment of female mosquitoes emerging from B vitamin-treated larvae. A transcriptomic analysis of male and female larvae identified some sex-specific regulated genes upon vitamin treatment. When treating germ-free larvae with individual B vitamins, we detected a specific toxic effect related to biotin (vitamin B7) exposure at high concentrations. We then provided germ-free larvae with varying biotin doses and showed that males are sensitive to biotin toxicity at a lower concentration than females. Gnotobiotic larvae exposed to controlled low bacterial counts or with bacteria characterized by slower growth, show a male-enriched adult population, suggesting that males require less bacteria-derived nutrients than females. These findings shed new light on sex-specific nutritional requirements and toxicity thresholds during the development of insect larvae, which impact the sex ratio of adults.

## Introduction

*Aedes aegypti* mosquitoes are vectors of pathogens responsible for various human diseases, notably including yellow fever or dengue fever. They are predominantly concentrated in tropical and neotropical regions, posing a significant threat to approximately 3.9 billion people who could potentially contract dengue ^1^. Mosquitoes are holometabolous insects, undergoing a complete metamorphic transformation from aquatic larvae to terrestrial adults. Consequently, several adult physiological traits, such as size and lifespan, heavily hinge on the quality of the insect larval development, specifically influenced by the nutritional status during this developmental stage.

Mosquitoes depend heavily on their microbiota for vital nutrients crucial for their larval development. In their natural habitat, mosquito larvae primarily feed on microorganisms, particulate organic matter, and detritus present in their breeding sites ^2^. In controlled laboratory rearing environments, the main food sources for larvae typically comprise fish food or commercial pet food designed for rodents, cats, or dogs. Mosquito larvae hatched from microbe-free eggs and maintained in sterile conditions are halted in their development when provided with a sterile conventional diet ^3^. However, their development can be rescued when a live microbiota is introduced or when provided nutrient-rich diet and kept in the dark, strongly implying that the microbiota plays a fundamental role in furnishing to its mosquito host essential nutrients that are not present in the diet ^3–6^. These essential nutrients encompass critical elements such as essential amino acids, nucleosides, and B vitamins, which are beyond the synthetic capabilities of most insects ^4,6^. The investigation of mosquito nutritional requirements and how the microbiota affects larval development could unveil crucial metabolic insights that might serve as the foundation for innovative vector control strategies.

To unravel the mechanisms behind the intricate interactions between mosquitoes and their microbiota, we previously developed a method that enables the generation of germ-free mosquitoes at virtually any stage of their development. Our approach consists in the colonisation of *Ae. aegypti* mosquitoes with an *Escherichia coli* strain auxotrophic for two amino acids, d-Alanine (D-Ala) and meso-diaminopimelic acid (*m*DAP), that are bacteria-specific and crucial for bacterial cell wall synthesis. When bacteria are supplied alongside D-Ala and *m*DAP to germ-free larvae, mosquito development is rescued. As soon as the larval rearing medium is changed to sterile water deprived of bacteria and of D-Ala and *m*DAP, bacterial growth is arrested and germ-free larvae are obtained. Using this method, we have been able to pinpoint folic acid as one of the critical metabolites furnished by the microbiota and required during mosquito larval development ^4^.

Vitamins of the B group are cofactors carrying on numerous cellular processes, encompassing essential functions in the electron transport chain (riboflavin, nicotinic acid), amino acid metabolism (pyridoxine, folic acid), lipid metabolism (biotin, riboflavin, nicotinic acid), and nucleic acid metabolism (folic acid, nicotinic acid)^7^. They are required in very low amounts, as they serve as coenzymes and are not consumed by the enzymes for which they act as cofactors ^8^. Insects depend on their food and microbiota to supply B vitamins as they lack the complete metabolic pathways to synthesize them ^9^. The insect model *Drosophila melanogaster* requires all B vitamins for optimal larval development ^10^ and similar necessities have been hypothesized for mosquitoes. As a matter of fact, B vitamins have been identified, along with sterols, aminoacids and nucelosides, as compounds in synthetic diets designed for rearing mosquito larvae in the absence of a microbiota ^11–13^. A more recent study has highlighted the critical roles played by amino acids and B vitamins in mosquito larval growth. Notably, riboflavin has been shown to be required throughout the entire larval development as it is light sensitive, and thiamine, pyridoxine, and folic acid were found to be specifically required by *Ae. aegypti* larvae to initiate pupation ^6^.

In our previous study, we found that folic acid provision partially rescued the development success of mosquito larvae deprived of their microbiota during late instars ^4^. Here, we tested the effect of supplying different B vitamin doses on *Ae. aegypti* larval development. Larvae cleared of their microbiota during their third instar and supplemented with increasing amounts of B vitamins did not show any improvement in their developmental success when compared to germ-free non-supplemented counterparts. Surprisingly though, we observed an impact on the sex-ratio of the resulting adults, with a significant enrichment of female mosquitoes emerging from B vitamin-treated larvae. We further investigate potential mechanisms involved in this sex-specific effect of B vitamins via transcriptomics, diet supplementation and bacterial monocolonisation. We show that during larval development, males require less bacteria-derived products, notably biotin, than females and that they are sensitive to lower doses of biotin, resulting in the observed sex ratio distortion.

## Results

### Vitamin supplementation affects mosquito sex-ratio in axenic conditions

We previously reported that *Ae. aegypti* larvae cleared of their microbiota at the third instar showed lower developmental success compared to conventionally reared larvae, with only 10–20% of individuals successfully completing the metamorphosis stage. The proportion of germ-free individuals reaching the adult stage increased to ∼50–60 % if folic acid (vitamin B9) was supplemented to the larval diet (0.25–1.25 mg/mL), suggesting an important role of the microbiota in providing this B vitamin to larvae ^4^. As the rescue was not complete, we wondered if other B vitamins potentially produced by the microbiota were involved in mosquito larval development and could further increase the proportion of larvae developing into adults. We produced germ-free third-instar larvae, following our transient colonization protocol ^4^: after egg sterilization, we mono-colonized first instar larvae with the *E. coli* HA416 strain, auxotrophic for D-Ala and *m*-DAP, in the presence of these amino acids, and starved larvae from D-Ala and *m*-DAP shortly after reaching the third instar to turn them germ-free. We supplemented these larvae with a solution containing six B vitamins (biotin, folic acid, nicotinic acid, pyridoxine, riboflavin, and thiamine) and choline (not classified as B vitamin anymore, but recognized as essential nutrient), at concentrations previously reported to support *Ae. aegypti* larval development in germ-free conditions (VIT1x ^13^). We did not observe any significant increase in the percentage of adult mosquitoes (Supplemental Figure S1, proportion of adults: *p* = 0.55, not significant (ns); see Supplemental Table S1 for details on statistics). Since the folic acid concentration supplemented in our previous study was 5–25 times more concentrated that the concentration used in ^13^, we decided to test the effect of four and eight times higher B vitamin concentrations (VIT4x and VIT8x) on larval development. Again, we did not observe an increase in the proportion of adult mosquitoes, rather a marginally significant decrease with VIT8x (Figure 1A, proportion of adults: *p* < 0.0001; AUX vs GF, VIT4x, or VIT8x: *p* < 0.0001; GF vs VIT4x: *p* = 0.73; GF vs VIT8x: *p* = 0.058; Supplemental Table S1). However, under germ-free conditions, the small number of mosquitoes that completed their development was due to an equal proportion (∼30%) of larvae either dying or being stalled at the larval stage. In contrast, the vitamin treatment significantly increased mortality rates, doubling them to approximately 60 % (Figure 1A, proportion of dead larvae: *p* < 0.0001; AUX vs GF, VIT4x, or VIT8x: *p* < 0.0001; GF vs VIT4x: *p* = 0.010; GF vs VIT8x: *p* < 0.0001; Supplemental Table S1). Interestingly, we saw a significant shift in the sex ratio of the fully developed mosquitoes, with more than 80 % of adult mosquitoes being females when larvae were treated with VIT4x and VIT8x solutions (Figure 1B, *p* < 0.0001; AUX or GF vs VIT4x: *p* = 0.0010; AUX vs VIT8x: *p* = 0.0037; GF vs VIT8x: *p* = 0.0019; Supplemental Table S1). This suggested a sex-specific effect of B vitamins on mosquito development.

**Figure 1.**
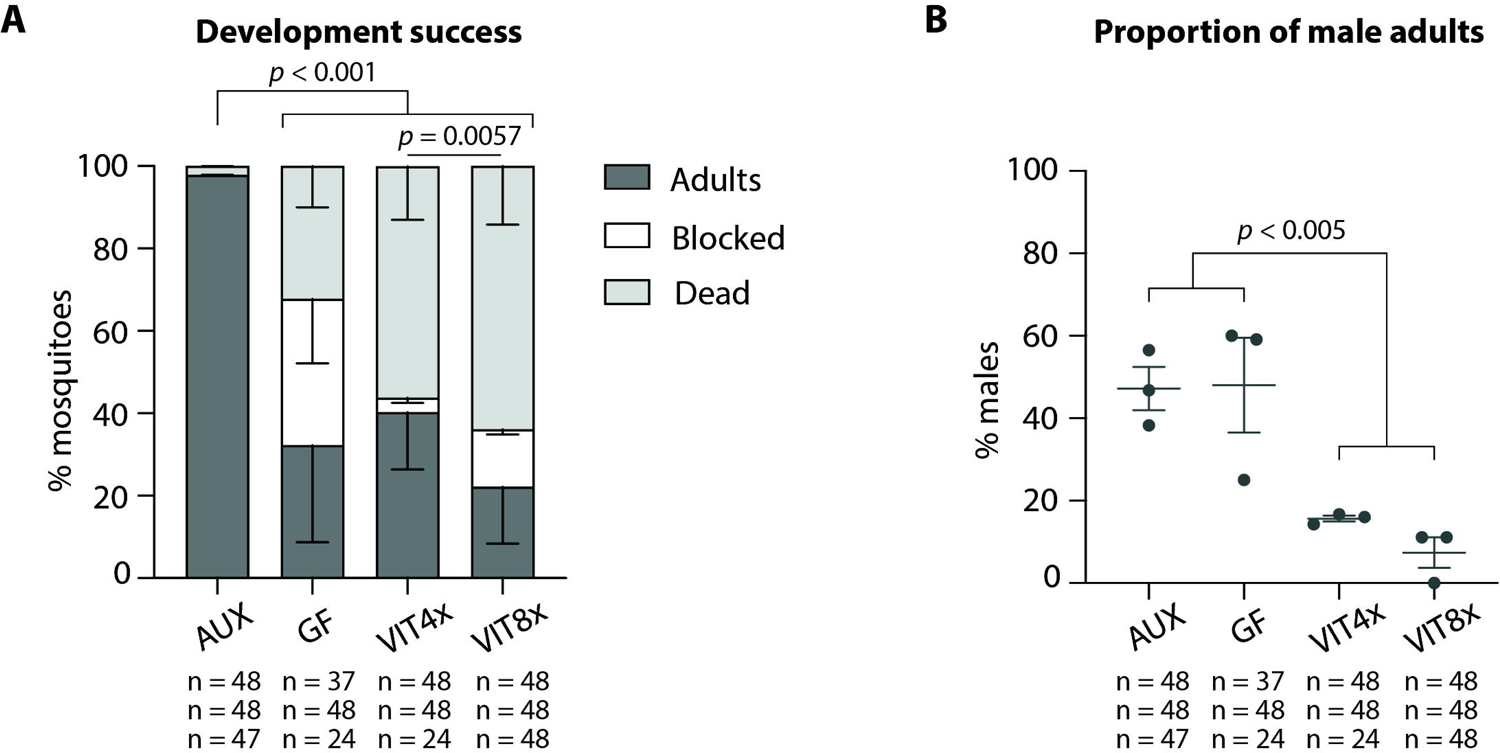
Development success and sex-ratio of mosquitoes exposed to B vitamins during larval development. (**A**) Proportion of individuals completing their development (dark grey), blocked at the larval stage (white), or dying during development (light grey) when initially reared with the auxotrophic *E. coli* strain (AUX), reversibly colonized at the beginning of the third instar and kept thereafter in germ-free conditions (GF) or supplemented with a 4x (VIT4x) or 8x (VIT8x) B vitamin solution. Bar charts represent the mean ± SEM of three independent replicates. (**B**) Proportion of male adult mosquitoes emerging from the larvae treated in (**A**). Numbers below graphs indicate the number of mosquitoes analysed per replicate and condition. See Supplemental Table S1 for detailed statistical information.

### Transcriptomic analysis of germ-free larvae treated with B vitamins

We had two alternative hypotheses to interpret the observed impact of B vitamins on sex ratio without significantly affecting overall development success. This effect can be attributed to a concomitant increase and decrease in the fitness of female and male larvae respectively upon vitamin supplementation, due to differential nutritional needs and to toxic effects of these compounds at the tested concentrations. Alternatively, some feminization mechanism may influence male trait maturation during larval development. To sort out which of the two possibilities prevailed, we conducted a transcriptomic study on male and female larvae cleared of their microbiota at the beginning of the third instar, kept in germ-free conditions (GF), and then supplemented with VIT8x until sampling, 3-6 h after the beginning of the fourth larval instar (Figure 2A; this transcriptomic study is detailed in Supplemental data and Supplemental Figures S2, S3). We observed on one hand that VIT 8x induced the heat-shock-protein related gene *AAEL017976* in males, which would lend credence to the toxicity hypothesis (Figure 2B). In parallel, it down-regulated *AAEL012340* and up-regulated *AAEL017067* in females, assigned to encode lipase 1 precursor and peritrophin-48-like, which also may point to a better development in females (Figure 2D, E). Conversely, we also observed a strong induction of *AAEL013606* in males, predicted to encode a SRY (Sex-determining region of Y chromosome)-like protein, which is a sex-determination factor and may alternatively point to a potential feminizing impact of B vitamins (Figure 2C). Hence, our transcriptomic analysis did not provide a clear conclusion on the way vitamins affected the sex-ratio.

**Figure 2.**
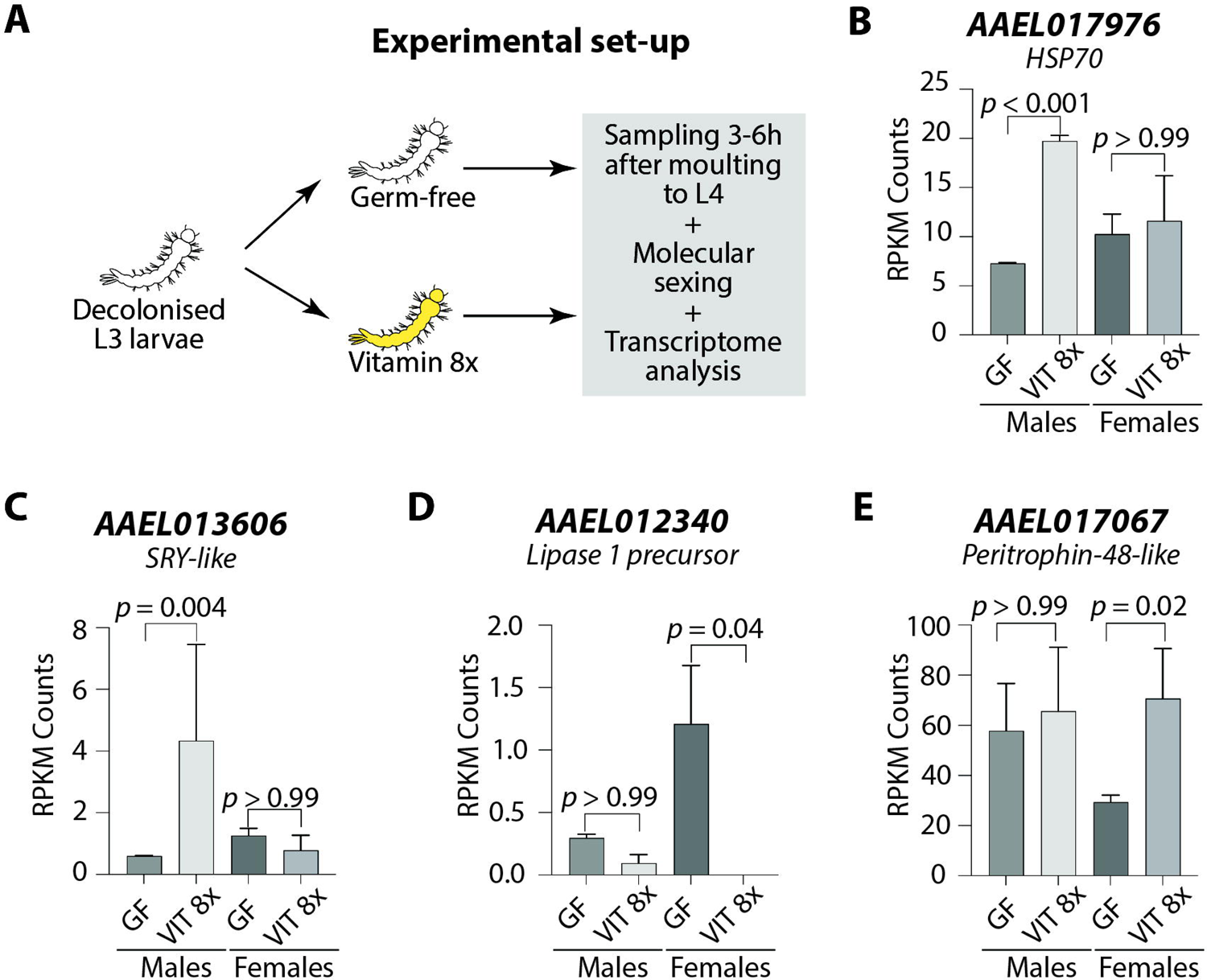
Transcriptomic analysis of male and female larvae exposed to B vitamins. (**A**) Experimental set-up: mosquito larvae reared in gnotobiotic conditions with the auxotrophic *E. coli* strain were decolonized at the beginning of the third instar (L3) to obtain germ-free larvae. Larvae were either kept in germ-free conditions or treated with 8x B vitamin solutions. After three to six hours since moulting to the fourth instar (L4), larvae were individually collected. DNA was extracted from individual larvae for sex assignment using the *Nix* gene, while RNA was extracted for transcriptome sequencing on male and female larvae. A more precise version of the experimental set up can be found in Supplemental Figure S2. (**B-E**) Reads Per Kilobase per Million mapped read (RPKM) values of representative genes differentially regulated with B vitamins in males (**B, C**) and females (**D, E**). Genes are named according to their Vectorbase ID.

### Effect of individual B vitamins on larval development and adult sex ratio

We reasoned that if vitamins had a toxic effect on males, we could detect it more clearly at a higher concentration. For this reason, we treated decolonised *Ae. aegypti* larvae with vitamins at high dose, 50x compared to the reference concentration ^13^, using single vitamins to narrow down which B vitamin was causing this sex-specific effect. We added a 16x dose of folic acid to all tested conditions to obtain enough adult mosquitoes in the germ-free control group and have a more reliable comparisons on adult mosquito sex-ratio. We observed a significant decrease in larval development success in folic acid and biotin treatments (Figure 3A, proportion of adults: *p* < 0.0001; GF vs folic acid: *p* = 0.015; GF vs biotin: *p* < 0.0001; all other comparisons *p* > 0.05, ns; Supplemental Table S1). In particular, while ∼90 % larvae developed into adults in the germ-free control, only ∼60% and ∼5 % mosquitoes reached the adult stage in the folic acid and biotin treatments, respectively. While the addition of folic acid induced a non-significant increase in the proportion of mosquitoes that were blocked in their larval development (from ∼2 % in the control to ∼ 20 %, Figure 3A, *p* = 0.0002; GF vs folic acid: *p* = 0.062, ns; Supplemental Table S1), biotin induced a strong mortality on larvae (from ∼10 % in the control to ∼70 %, Figure 3A, *p* < 0.0001; GF vs biotin: *p* < 0.0001; Supplemental Table S1). The sex-ratio of the resulting adults was not significantly affected by any vitamin supplementation, although no viable male mosquito developed from biotin-treated larvae (Figure 3B, *p* = 0.73, ns; Supplemental Table S1). This low statistical effect was probably due to the small number of biotin-treated larvae that could complete their development (adult mosquitoes per replicate (sex): 1/24 (non identified); 1/24 (female); 3/24 (females)), suggesting that the biotin 50x dose was generally toxic to both sexes. However, we could not determine if the absence of fully developed males after biotin supplementation was due to a male-killing effect or to the feminization of phenotypic characters in male mosquitoes.

**Figure 3.**
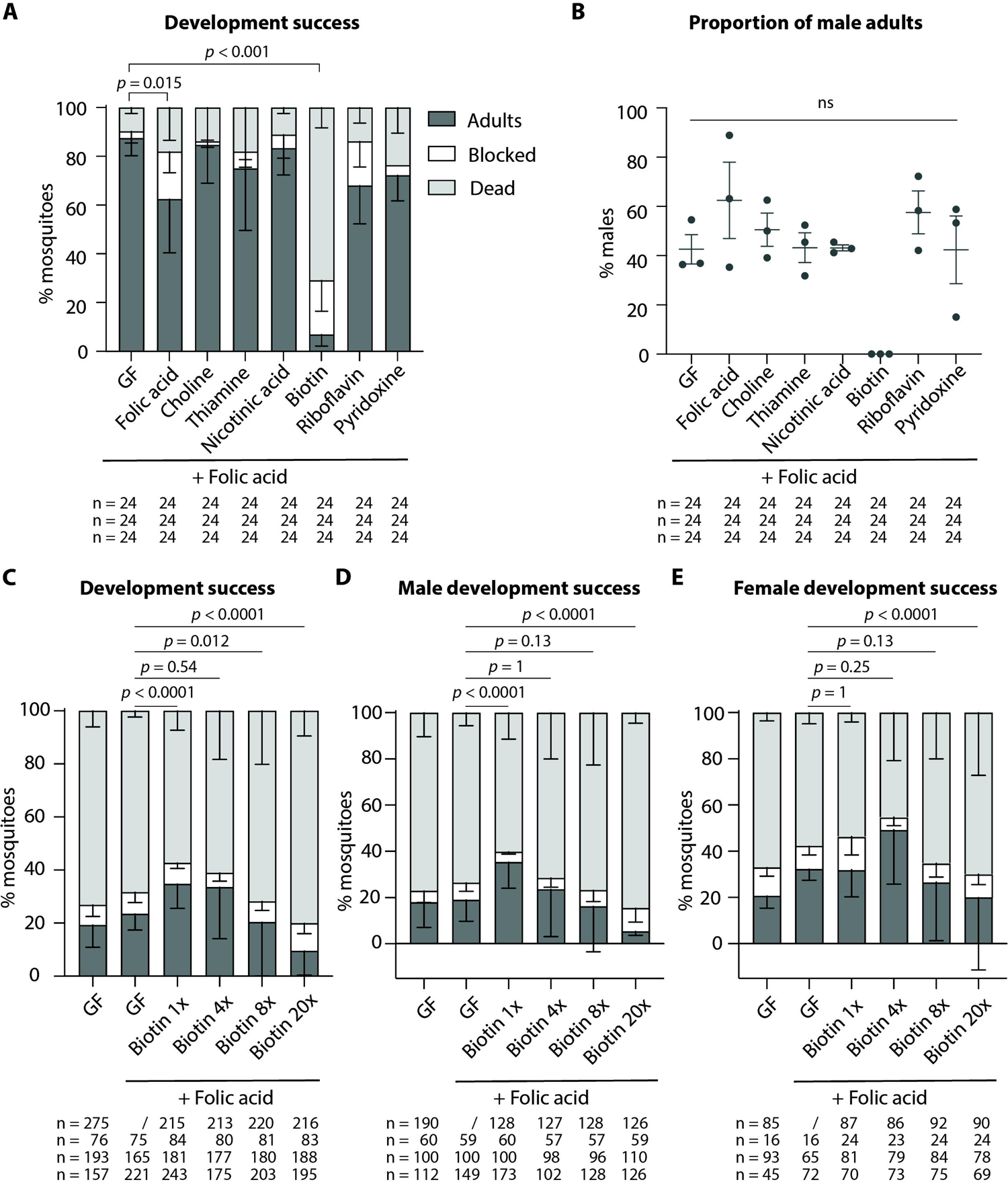
Development success of male and female mosquito larvae exposed to high doses of single B vitamins or to increasing doses of biotin. (**A**) Proportion of mosquitoes completing their development (dark grey), blocked at the larval stage (white), or dying (light grey) when reversibly colonized at the beginning of the third instar and kept in germ-free conditions (GF) or supplemented with a 50x solution of folic acid, choline, thiamine, nicotinic acid, biotin, riboflavin, or pyridoxine. A 16x folic acid solution was also added to all tested conditions. Bar charts represent the mean ± SEM of three independent replicates except for riboflavin (two replicates). (**B**) Proportion of male adult mosquitoes emerging from the larvae treated in (**A**). (**C-E**) Proportion of mosquitoes (Aaeg-M strain) completing their development (dark grey), blocked at the larval stage (white), or dying (light grey) when reversibly colonized at the beginning of the third instar and kept in germ-free conditions (GF) or provided with a 1x, 4x, 8x, or 20x biotin solution. A 16x folic acid solution was also added to all tested conditions. Bar charts represent the mean ± SEM of four independent replicates except for GF+folic acid (three replicates). Data in (**C**) are not sorted by sex, while data in (**D**) and (**E**) represent results for male and female mosquitoes, respectively. Numbers below graphs indicate the number of mosquitoes analysed per replicate and condition. See Supplemental Table S1 for detailed statistical information.

### Biotin requirements and toxicity are sex-specific

The experimental set up used in previous experiments could not distinguish between biotin-related male toxicity or feminization of male mosquitoes. In fact, our visual analysis of fully-developed mosquitoes indicated that adult mosquitoes displayed female-specific phenotypic characteristics, but this analysis did not yield information on the genotype of those mosquitoes and of those that did not reach adulthood. We took advantage of a recently established *Ae. aegypti* genetic sexing strain (Aaeg-M) characterized by the insertion of the *eGFP* transgene in the male-specific M locus ^14^. This allowed us to sort first instar larvae by sex right after egg sterilization and perform the full experiment on male and female larvae in parallel. The effect of four increasing biotin concentrations (1x, 4x, 8x and 20x) was tested on decolonized third instar larvae. As done previously, we added 16x folic acid to all tested solutions to increase the number of adult mosquitoes.

When analysing the global results independently from the mosquito sex, we observed that the addition of folic acid alone did not significantly increase the proportion of larvae completing their development to adulthood in the Aaeg-M strain, with ∼20 % of fully developed adults in both germ-free controls, with or without folic acid (Figure 3C, *p* < 0.0001; GF vs GF+folic acid: *p* = 1, ns; Supplemental Table S1). The proportion of fully developed mosquitoes at 1x biotin concentration increased to ∼30 %, while it decreased to ∼20 % and ∼10 % in 8x and 20x biotin treatments, respectively (Figure 3C, see Supplemental Table S1 for individual comparisons). In all treatments, the variation in the proportion of adult mosquitoes was due to an equivalent change in the percentage of dead mosquitoes rather than of stunted larvae.

Data sorted by sex indicated a differential biotin requirement for male and female mosquitoes: while the biotin 1x concentration was the best one to support the development of males (∼ 30 %, Figure 3, *p* < 0.0001, see Supplemental Table S1 for individual comparisons), the 4x concentration resulted in the highest development success in females (∼ 50 %, Figure 3E, *p* < 0.0001, see Supplemental Table S1 for individual comparisons). In females, the effect of the addition of biotin 4x was not statistically different from germ-free conditions supplemented with folic acid (*p* = 0.25, ns; Supplemental Table S1), but the addition of biotin 4x significantly improved the proportion of developed mosquitoes in the absence of folic acid (*p* = 0.0001; Supplemental Table S1). As discussed below, the sole addition of folic acid did not significantly improve the development of germ-free larvae in these conditions (*p* = 1, ns; Supplemental Table S1). To confirm that the toxic effect was specific for biotin and not a combinatory effect between biotin and folic acid, we tested these vitamins separately. Indeed, a 50x treatment with biotin alone significantly reduced the proportion of developed mosquitoes, in both males and females (Supplemental Figure S4A-C).

When comparing male and female mosquitoes in their developmental rates to pupa (i.e., the proportion of larvae starting metamorphosis by day), we observed similar proportions of mosquitoes starting metamorphosis in all conditions, with a higher percentage of males going through pupation in germ-free (*p* < 0.0001), biotin 1x (*p* < 0.0001) and 8x (*p* = 0.02) treatments (Supplemental Figure S5A; Supplemental Table S1). However, a significant proportion of these pupae were not able to complete the developmental stage and died before emergence (Supplemental Figure S5B). Interestingly, the addition of 16x folic acid alone to germ-free larvae differentially affected the pupation rates of male and female mosquitoes, suggesting that sex-dependent requirements are not exclusive for biotin (*p* = 0.0005; Supplemental Figure S5B; Supplemental Table S1). Taken together these data suggest that the clearance of the microbiota during the third instar impacts mosquito development at the metamorphosis stage, and that the addition of biotin at a 1x concentration improves male development while a 4x concentration already causes toxicity; in females, the optimal concentration for development is 4x. Since toxicity might be generally correlated to body size, we tested whether male larvae had a significant lower mass or size compared to female larvae, similarly to what observed in adult mosquitoes. Third instar larvae did not show differences between sexes when measuring total dry mass or larval length (dry mass: *p* = 0.13; body length: *p* = 0.54, Supplemental Figure S6A-B). However, male larvae had significant smaller head capsules (a proxy of body size) compared to females (*p* = 0.01, Supplemental Figure S6A). Given that the weight difference between males and females is less than 6 % and the difference in head capsule size is less than 2 %, we conclude that the variations in biotin toxicity are likely attributable to other sex-specific factors beyond size.

For each experiment, adult mosquitoes were visually inspected to confirm that their genotype (GFP fluorescence status in L1) matched their adult phenotype. In parallel, an experiment with New Orleans mosquitoes was performed only on decolonised larvae kept germ-free or supplemented with 4x or 8x biotin. To compare mosquito phenotype and genotype, emerged mosquitoes were visually analysed to confirm their sex. Their DNA was extracted and subjected to dual PCR on *Nix* and *Actin*. PCR amplicons were analysed using a capillary electrophoresis system that allowed to automatically detect gene-specific signals. Among the analysed mosquitoes (GF: n = 70, biotin 4x: n = 65, biotin 8x: n = 59), none showed discordant results between genotype and phenotype, further confirming that biotin had a toxic effect but did not induce any male feminization.

### Males require less bacteria-derived metabolites than females for development

Based on these results, we hypothesized that males required lower quantities of bacteria-derived metabolites than females. To test this, we provided larvae with a suspension of auxotrophic bacteria approximately 10^6^ times more diluted than the standard solution used to support conventional mosquito development. This setup required active bacterial growth during larval development, leading to a greater dependency on the amino acids *m*-DAP and D-Ala for bacterial proliferation and larval development. Thus, a *m*-DAP and D-Ala solution at a conventional (1x) or 100-fold-diluted (0.01x) concentration were provided every three days until pupa appearance. This allowed to maintain a proliferating bacterial population while ensuring that bacterial loads were 2.2-73 fold lower when provided the diluted solution (Figure 4A, 1x vs 0.01x: day 1, day 2, day 3, day 6, day 7, day 10: *p* < 0.0001; day 8: *p* = 0.013; day 9: *p =* 0.0014; Supplemental Table S1). Larvae received the same amount of larval food in both conditions. This reduction in bacterial counts in larva water led to a lower development success (76 % to 33 %, *p* < 0.0001, Figure 4B). Development was also slower (11.4 days to 12.4 days: *p =* 0.012; Figure 4D; Supplemental Table S1). While females generally develop slower than males, this did not explain slower development as larvae that developed in 15 days or more in the 0.01x condition were only males. As expected, sex ratio was also distorted towards a higher proportion of males with lower amounts of bacteria (Figure 4C; 59% to 89 %, *p* = 0.043; Supplemental Table S1). When doing a 10-fold dilution (0.1x), we observed similar trends, albeit with non-significant differences on sex ratio, partly because less replicates were performed (Supplemental Figure S7A–D; % adults: *p* = 0.0061; sex: *p* = 0.32; time: *p* = 0.012; Supplemental Table S1).

**Figure 4.**
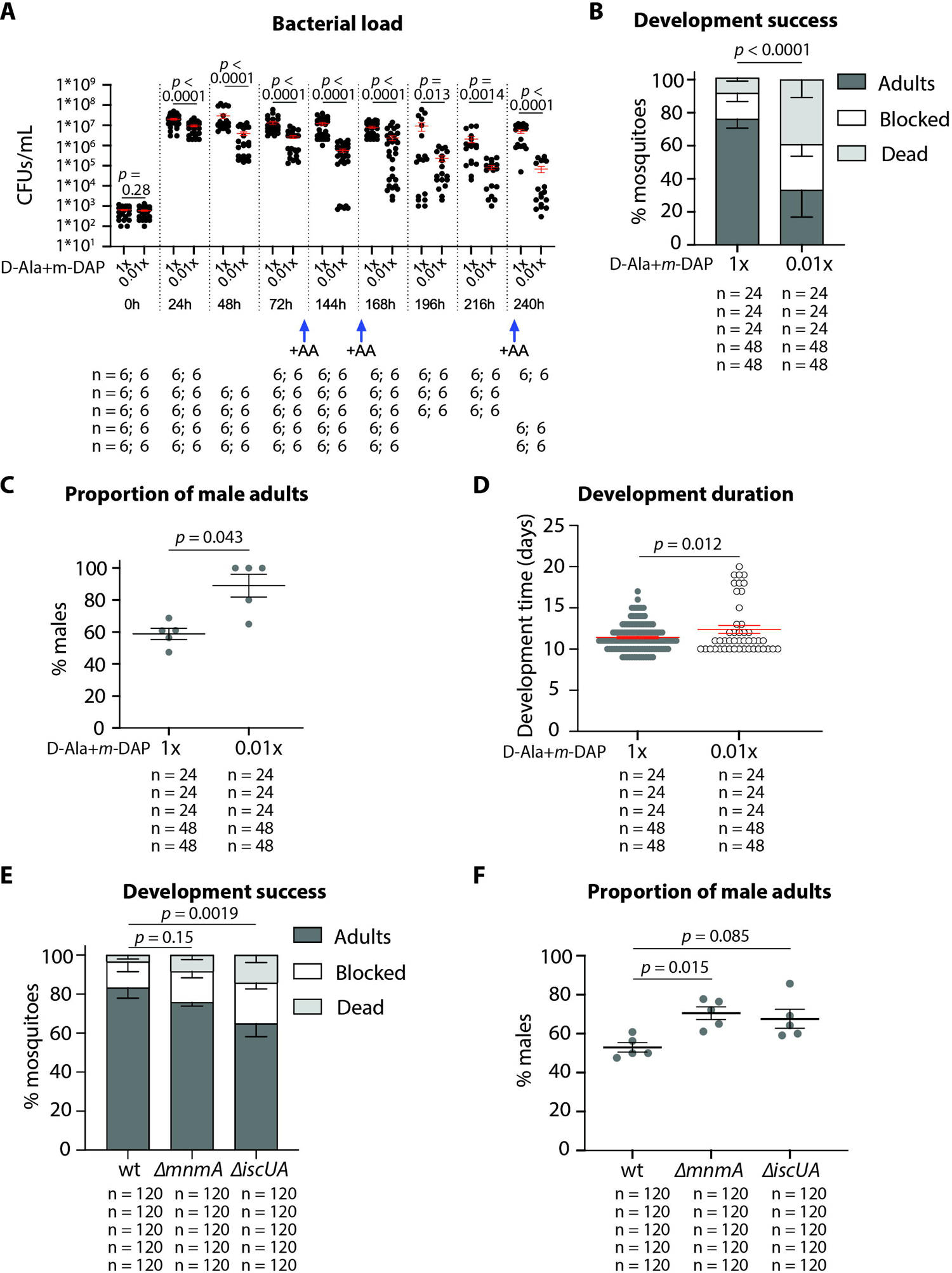
Development success and sex ratio of larvae with decreased support in bacterial metabolites. In **A-D**, auxotrophic *E. coli*-colonized larvae were treated with 1x or 0.01x concentrations of D-Ala and *m*-DAP to slow bacterial development. (**A**) Number of colony forming units (CFU)/mL in wells containing individual larvae. Each dot represents a well, 6 wells/time point/replicate were tested in at least 3 replicates. Time-points indicate the time after bacteria were added to germ-free larvae. (**B**) Proportion of mosquitoes reaching adulthood (dark grey), blocked in development (white) or dead (light grey) when auxotrophic *E. coli*-colonized larvae were supplemented with standard (1x) or diluted (0.01x) D-Ala and *m*-DAP concentrations. (**C**) Proportion of males amongst the adults and (**D**) number of days until reaching adulthood when auxotrophic *E. coli*-colonized larvae were supplemented with standard (1x) or diluted (0.01x) D-Ala and *m*-DAP concentrations. In **E-F**, larvae were colonised with wt *E. coli*, the growth deficient mutant Δ*mnmA* or the biotin-deficient Δ*iscUA* mutant. (**E**) Proportion of mosquitoes reaching adulthood (dark grey), blocked in development (white) or dead (light grey) when larvae were colonised with different *E. coli* mutants. (**F**) Proportion of males amongst the adults when larvae were colonised with different *E. coli* mutants. Numbers below graphs indicate the number of wells/mosquitoes analysed per replicate and condition.

We then wished to test whether such effects were specific to the provision of biotin. The use of a Biotin synthase (BioB) auxotrophic mutant was impossible in the experiment condition set-up because it would require the addition of biotin for its growth. We thus relied on a Δ*iscUA* mutant that is deficient in the main Fe-S cluster biogenesis pathway, hence predicted to be defective in the maturation of Fe-S proteins including BioB. BioB lacking this cluster was indeed reported to shows stability defect ^15^. However, our own unpublished results show that the Δ*iscUA* mutant is not auxotroph for biotin because the second Fe-S biogenesis system SUF is taking over and providing some level of maturation of BioB. We observed that Δ*iscUA* has a significantly lower ability to support larval development to adulthood than the control (65 % respect to 83 %: *p* = 0.0019; Figure 4E; Supplemental Table S1) and that it supported the development of a slightly higher proportion of males, albeit marginally significantly (*p* = 0.085; Figure 4F; Supplemental Table S1). This result supported the view that biotin level lowering bears consequence on larval growth and sex ratio. Another possibility is that the reduced efficiency of the Δ*iscUA* mutant in supporting larvae development may be due to its slower growth rate and compromised fitness. Therefore, to test this last possibility, we used the Δ*mnmA E. coli* mutant known to show slow growth rate ^16^ (Supplemental Figure S8). The gene *mnmA* encodes an enzyme required for thiolation of a subset of tRNAs. Larvae fed with Δ*mnmA* had an intermediate development success (*p* = 0.15; Figure 4E; Supplemental Table S1) and a significant increase in proportion of males among adults (*p* = 0.015; Figure 4F; Supplemental Table S1). Altogether these results indicated that microbial growth, and presumably associated richness in metabolite production, rather than specifically biotin level, is important for sex ratio.

## Discussion

The mosquito microbiota provides essential nutrients to its host. Among these nutrients, B vitamins have been shown to be required by mosquitoes to complete their larval development ^4,6,13^. Here, we found that the larval B vitamins requirements, especially biotin, are sex-specific: males require less B vitamins than females for a successful development. Together, the differences in nutritional requirements and in thresholds of toxicity led to a sex-ratio distortion after supplementing diet of germ-free *Ae. aegypti* larvae with increasing doses of B vitamins. Specifically, there was a notable reduction in the proportion of male mosquitoes emerging from larvae that had been subjected to vitamin supplementation. We conclude that males require less bacterial-derived metabolites, including biotin, for development.

Evolutionarily speaking, favouring male development when nutritional conditions are scarce may be a way to maintain a progeny that will be successful if it spreads and finds mating partners elsewhere, while when nutritional conditions are optimal, having a high development success of both sexes may allow an efficient local colonization. While our experiments were performed with individuals kept separately, the impact on sex ratio may be exacerbated in a population as male and female mosquitoes have different developmental dynamics at the larval stage, where females emerge later than males. In line with this, female adult body sizes are impacted more significantly than males by the composition of larval diet ^17^. Female development may be longer because mosquito larvae must reach a critical mass to successfully commence metamorphosis, and this mass is higher for females than males, both at the larval and adult stages ^18^. In species of scarab beetles where males are larger than females, instantaneous growth rate is not different between sexes but optimal growth lasts longer in males ^19^. Contrary to what we observe in mosquitoes, total development time is however not necessarily correlated with size in these beetles ^19^. Similarly in *Drosophila*, females are larger than males, yet they develop on average 4 hours earlier than males, indicating that final mass and development timing are not necessarily positively correlated. During fly metamorphosis, this protogyny phenotype is genetically controlled by *Sex lethal*, the master sex switch gene ^20^.

Diet composition at the larval stage has already been shown to affect the sex ratio of the adult mosquito population, with higher proportions of males emerging in starvation or suboptimal feeding conditions ^21^. Our data now specifies that males and females have different requirements in bacteria-derived metabolites, notably biotin, while the amount of provided food did not vary in our experiments. Females may also need more energy storage than males for egg production, as egg production after the first blood meal has been found to be strongly correlated with richness of the larval food ^22^.

The observed male-killing effect appears to be a specific outcome resulting from exposure to one or a combination of B vitamins when exceeding a concentration threshold. We observed that the optimal concentration of biotin for the development of germ-free female larvae was four times higher than what had been previously reported as sufficient to sustain mosquito larval development in germ-free conditions ^13^, while this concentration was already toxic to males. Depending on the vitamin, our 4x and 8x concentrations were also 2–160-fold higher than the concentrations used more recently to formulate a chemically defined medium for germ-free *Ae. aegypti* larvae ^6^. These two studies identified osmotic pressure as a critical limiting factor for artificial diets, describing a rapid larval mortality when the rearing medium contained either 11.6 g/L ^13^ or 113.7 g/L ^6^ of amino acids. The osmotic pressure of these solutions is ∼25 or ∼256 times higher than that of the 8x vitamin B solution we tested. Therefore, it is unlikely that the elevated mortality observed in our experiments could be due to high solute concentration. Yet, our transcriptomic analysis revealed the upregulation of a chloride channel encoding gene in both male and female larvae treated with the 8x vitamin solution, which may reflect a response to an increase in solute concentrations.

To investigate the potential mechanisms behind the male-specific toxic effect of B vitamins, we studied the transcriptomes of germ-free male and female larvae exposed to vitamin solutions. We encountered several problems in the process of sample sex assignment via PCR and RNA degradation, which led us to exclude one replicate and the entire vitamin 1x condition. This resulted in a relatively low number of genes that exhibited sex-specific regulation when compared to a prior transcriptomic study ^23^. With the same sex assignment technique, Matthews *et al*’s reported approximately 3400 differentially regulated genes between male and female fourth instar larvae. Between 35 % and 40 % of the genes identified in our transcriptomic study overlapped with those identified by Matthews *et al*, suggesting a reasonable degree of consistency with our data, considering that our larvae were germ-free. Most significantly, *Nix* was specifically enriched in male transcriptomes. These observations suggest that while we likely missed many regulated genes, those identified in our study are reliably true positives. The analysis of genes differentially regulated in vitamin 8x conditions identified different sets of genes in male and female larvae. Notably, the upregulation of genes linked to stress response, DNA binding, development or sexual reproduction was specific to male larvae. This observation suggests that the administration of high doses of vitamins interferes with critical cellular processes, as indicated by the upregulation of a heat-shock protein coding gene. The vitamin-induced stress response might explain the observed reduction in developmental rates among male larvae. However, it remains challenging to draw definitive conclusions regarding the precise mechanisms that underlie these effects.

When we administered individual B vitamins at elevated doses (50x), we observed that biotin, and to a lesser extent, folic acid, had lethal effects on germ-free larvae. However, only biotin was found to have an impact on sex ratio at this dose. Using the Aaeg-M GSS strain, we were able to distinguish the developmental success of male and female larvae independently and to validate the absence of a feminizing effect caused by biotin. To maintain consistent experimental conditions, we added folic acid to all tested conditions. Surprisingly, this addition in the absence of biotin did not significantly affect the proportion of fully developed adults in germ-free conditions, in contrast to what we typically observe (^4^, Figure 3A). This discrepancy could potentially be explained by differences in the mosquito strains used in the two sets of experiments (New Orleans and Aaeg-M GSS, which was created from mosquitoes sourced in Thailand ^14^). However, we think that the variations in the timing between the two types of experiments were responsible for the different outcomes of folate supplementation. Due to the large number of larvae and to diet autofluorescence, first instar larvae of the Aaeg-M GSS strain had to be starved for 24 h while assigning the sex via GFP fluorescence before adding bacteria and food. Hence, larvae were initially in a deprived metabolic status which may have carry-over effects on later development. We hypothesize that folic acid alone is not sufficient to complement their requirements. Furthermore, experiments showed a noteworthy degree of variability of development success between replicates. While experimental conditions were extremely controlled, replicates were conducted with different batches of eggs produced in conventional conditions. This variability underscores the influence of various factors such as the maternal gonotrophic cycle or the age of the eggs on the outcomes of larval development. The quality of the blood on which mothers have fed and the quantity of eggs laid by each female could potentially influence the amounts of vitamins in embryos, consistent with observations that arthropod development is influenced by mother’s age and diet^7,24,25^.

Using the GFP-sexing strain, we showed that male and female mosquito larvae cleared of their microbiota exhibit different biotin requirements for their development (0.5 µg/mL for males and 2 µg/mL for females). These different biotin requirements and observed toxicity at higher doses might correlate with larval size, which tends to be larger in females than in males during the larval stage. Importantly, both sexes show heightened mortality rates during metamorphosis when exposed to higher biotin concentrations. Although more than 50 % of the larvae successfully transitioned into pupae under all tested conditions, only a range of 10 to 40 % emerged as fully developed adults (see Supplemental Figure S5A). This observation suggests that exposure to elevated biotin levels during the third and fourth instars predominantly impacts pupae. Biotin has been found to be essential for intestinal stem cell mitosis in *Drosophila* ^26^, and is more generally involved in cell cycle progression ^27^, suggesting its importance during life stages with high cell proliferation rates. During metamorphosis, tissue remodelling increases cell proliferation, hence the number of biotin-responsive cells may be relatively high. Elevated biotin doses may specifically deregulate cell cycle at this stage and explain why we observe high pupal mortality. In line with our observation, previous studies underscored the essential role of biotin in *Ae. aegypti* larval development, but they also showed that high biotin concentrations induced mortality in pupae when larvae were reared in both axenic or conventional conditions ^28,29^. Some degree of biotin toxicity has been observed in pupae of the flour beetle *Tribolium confusum* as well ^30^. Furthermore, high biotin concentrations that still allowed *Ae. aegypti* development were reported to reduce adult fertility; they notably caused follicle degeneration in females during the post-ovipositional period^29^. This effect of biotin on fertility seems to be conserved in other insect species such as the Mexican fruit fly *Anastrepha ludens* ^31^, the house fly *Musca domestica* ^32^ and the beetle *Dermestes maculatus*^33^. In these insects, females tended to lay fewer eggs when fed biotin-rich diets and eggs showed lower hatching rates. In *D. maculatus* the impact of biotin on embryogenesis appeared to be related to the binding of this vitamin to insoluble yolk proteins, resulting in reduced amino acid availability within the embryo ^34^.

Mosquitoes are incapable of synthesising B vitamins, thus relying on their diet and microbiota for the essential provision of these metabolites ^4,6–8^. Notably, germ-free mosquito larvae can only develop if a rich diet is provided and if measures are taken to protect vitamins from light-induced degradation ^5,6^. Several bacterial strains including *E. coli* are capable of rescuing mosquito development when introduced to germ-free larvae, while others such as *Microbacterium* are not ^3,4^. This correlates with genomic data showing that *E. coli* possess complete metabolic pathways for B vitamin synthesis ^7^, while *Microbacterium* does not ^3,35^. Our findings regarding biotin toxicity and previous data indicating biotin impact on insect fertility even raise the possibility that bacteria might interfere with mosquito sex ratio or, more generally, with mosquito physiology, by delivering high concentrations of some B vitamins. Intriguingly, genomic analyses of *Wolbachia* genomes have identified a riboflavin transporter gene to be associated with cytoplasmic incompatibility ^36^. Although vitamin concentrations used in our experiments differ significantly from what mosquito would encounter in natural conditions, our findings underscore the critical need to explore bacterial-induced mechanisms that involve B vitamins because they influence the mosquito host larval development, toxicity, fertility, and the alteration of the sex-ratio.

## Materials and Methods

### Mosquitoes and bacteria

*Escherichia coli* HA 416 was grown in lysogeny broth (LB) supplemented with 50 µg/mL kanamycin, 50 µg/mL *m*-DAP and 200 µg/mL D-Ala. For all experiments, *E. coli* cultures were inoculated from single fresh colonies in liquid LB with appropriate supplementation and incubated at 30 °C, shaking at 150 rpm for 16 h. *E. coli* FBE051 (wt), FBE356 (Δ*iscUA*) and FBE584 (Δ*mnmA*), used for Figure 4D-E, were cultured in the same conditions in LB with (Δ*iscUA* and Δ*mnmA*) or without (wt) 50 µg/mL kanamycin supplementation.

*Aedes aegypti* mosquitoes belonged either to the New Orleans strain or to the Aaeg-M genetic sexing strain (GSS, ^14^). Both colonies were maintained under standard insectary conditions at 28-30°C on a 12:12 h light/dark cycle. Mosquitoes are routinely blood-fed either on beef blood provided by the local slaughterhouse (Abattoir Territorial de Rémire Montjoly) or on anesthetized mice. The protocol of blood feeding on mice has been validated by the French Direction générale de la recherche et de l’innovation, ethical board # 089, under the agreement # 973021. Gnotobiotic mosquitoes were maintained in a climatic chamber at 80 % relative humidity on a 12:12 h light/dark 28 °C/25 °C cycle.

### Vitamin solutions

B vitamins (Sigma-Aldrich) were dissolved in the appropriate solvent at the concentration indicated in Table 1. Stock solutions were adjusted to pH 7.4-8.0 and filtered through a 0.22 µM membrane filter and stored at −20 °C until use.

**Table 1.** List of the B vitamins tested with their reference and stock concentrations, and the solvent used for stock solutions.

| Vitamin | Reference concentration <sup>13</sup> | Solvent | Stock concentration |
| --- | --- | --- | --- |
| Biotin (B7) | 0.5 µg/mL | NaOH 0.1 M | 0.4 mg/mL |
| Choline | 25 µg/mL | Water | 50 mg/mL |
| Folic Acid (B9) | 30 µg/mL | NaOH 0.1 M | 50 mg/mL |
| Nicotinic Acid (B3) | 10 µg/mL | NaOH 0.1 M | 50 mg/mL |
| Pyridoxine (B6) | 10 µg/mL | Water | 50 mg/mL |
| Riboflavin (B2) | 20 µg/mL | NaOH 0.1 M | 10 mg/mL |
| Thiamine (B1) | 10 µg/mL | Water | 50 mg/mL |

### Generation of gnotobiotic larvae

Germ-free larvae were obtained as previously described ^4^. Briefly, eggs were placed on top of a filtration unit and surface sterilised by subsequent washes in 70 % ethanol for 5 min, 1 % bleach for 5 min, and 70 % ethanol for 5 min. After rinsing three times with sterile water, eggs were transferred to a sterile 25-cm^2^ cell-culture flask filled with ∼20 mL of sterile water. Approximately 20-30 eggs were inoculated in 3 mL of liquid LB and incubated for 48 h at 30 °C shaking at 150-200 rpm to confirm sterility.

The following day (day two), a 16 h culture of auxotrophic *E. coli* was centrifuged, and the bacterial pellet was resuspended in 5 times the initial culture volume of sterile water supplemented with *m*-DAP (12.5 µg/mL) and D-Ala (50 µg/mL). Larvae were individually placed in 24-well plates together with 2 mL of bacterial suspension and ∼50 µL of autoclaved TetraMin Baby fish food suspension. Larvae were kept in a climatic chamber at 80 % RH with 12:12 h light/dark 28 °C/25 °C cycle.

To achieve bacterial decolonisation, larvae that moulted to the third instar in a time window of 5 h during day 4 were washed in sterile water and individually transferred in a 24-well plate filled with 1.5 mL of sterile medium and 50 µL of sterile fish food in each well. Larvae obtained with this method were shown to be largely germ-free after 12 h since transfer ^4^. Routine checks were performed to detect bacterial contamination. Specifically, the water used to rinse larvae was plated in LB plates to verify the absence of contaminating bacteria, and in LB plates supplemented with *m*-DAP, D-Ala, and kanamycin, to check for the carryover of auxotrophic *E. coli*. Additionally, for all experiments monitoring larval growth, replicates showing high development rate were tested for bacteria contamination in later time-points and excluded if bacterial growth was detected.

The Aaeg-M GSS required a sex-sorting step prior to being transiently colonised by auxotrophic *E. coli*. This strain carries an *eGFP* transgene in the male-specific M locus, thus only male mosquitoes express GFP. Axenic first instar larvae were individually placed in 96-well plates and checked for green fluorescence under an EVOS FL Auto System (Thermo Scientific). After sex-sorting, larvae were individually placed in 24-well plates. Bacteria were added the following day (day three) therefore the experiment schedule was shifted by one day.

For experiments shown in Figure 4 and Supplemental Figure S7, bacterial suspensions were diluted in sterile water to 100 CFU/mL (i.e., approximately 10^6^ times more diluted than in the normal setup). In Figure 4A-D and Supplemental Figure S7, amino acids were provided at a conventional concentration (1x, *m*-DAP – 12.5 µg/mL and D-Ala – 50 µg/mL) or after 1:10 or 1:100 dilution in sterile water. For each condition, the initial amino acid concentration was provided again every three days until pupae appeared.

### Vitamin treatment of axenic larvae

At the time of transfer, germ-free third instar larvae were randomly assigned to the different treatments and kept individually in 24-well plates. The larval medium consisted either of sterile water alone (germ-free condition) or sterile water supplemented with a single vitamin or with a mix of B vitamins (see Table 1 for reference B vitamin concentration). Sterile fish food was supplied to each larva. The 24-well plates were placed in a climatic chamber at 80 % RH with 12:12 h light/dark 28 °C/25 °C cycle into plastic boxes covered with aluminium foil to prevent B vitamins from light-degradation ^6^.

### Analysis of axenic larvae developmental success in different vitamin conditions

Larvae treated with different B vitamin conditions were observed daily to track their developmental success. For each larva, the following parameters were recorded: day of moulting to the fourth instar, day of pupation, day of adult emergence, sex; in case of failed development to adult, the day of death was recorded. Larvae were monitored for 14 days after transfer in the germ-free medium.

If at the end of this period larvae were still alive but did not undergo metamorphosis, they were marked as “blocked”.

### Transcriptomic analysis

All methodological details about transcriptomics are described in Supplemental material.

### Dry mass, larval length and head capsule width measurements

Gnotobiotic Aaeg-M GSS larvae were obtained as described above and monitored for their development. Larvae were sampled within 5 hours after moulting to the third instar, sorted by sex checking green fluorescence, and either used to quantify body dry mass (n = 50 per sex/replicate) or to measure body length and head capsule width (n = 20 per sex/replicate). Larvae collected to quantify dry mass were pooled, dried for a total of 2 h in a SpeedVac Concentrator plus (Eppendorf) using the following program: 90 min of vacuum, 2 h of pause, 30 min of vacuum. Body length and head capsule width were measured on larvae immobilized in 60 % ethanol using a Leica M165 C stereo microscope (Leica Microsystems). Larval body length was measured as the distance between the anterior border of the head and the posterior border of the last abdominal segment, excluding the siphon. Head capsule was measured as the distance between the root of the two antennae. Three independent replicates were performed.

### CFU quantification

For each condition, 20 µL of breeding water from 6 different wells were sampled and each sample was transferred into 200 µL of sterile water. For each time point, sampling was randomly carried out among the wells whose larvae were at the most advanced stage. For CFU counts, bacterial suspensions were serially diluted in sterile water and 10 µL was spotted on LB agar plates, incubated at 30 °C and counted 24 h later.

### Growth curves

Bacteria were grown overnight in LB medium at 30 °C then diluted in LB at 1:50. Growth curves were produced by quantifying optical density at a wavelength of 600 nm every 30 min during 24 hours in a FLUOStar Omega plate reader (BMG Labtech). Plates were shaken before each measurement. All the conditions were performed in triplicates (i.e. starting from three distinct colonies) and technical duplicates.

### Statistical analyses

Graphs were created with GraphPad Prism (version 10.0.2). Statistical analyses were performed with generalised linear mixed models (GLMM) using the lme4, lmerTest and lsmeans packages in R (version 4.3.0). For categorical data (development success, sex ratio) the replicate was set as a random effect and an ANOVA was performed on a logistic regression (glmer). For quantitative data (CFU counts, duration of development, larval length measurements) an ANOVA was performed on a linear regression (lmer). To compare larval body weight a paired t-test was performed. For developmental rate analyses, a Log-rank (Mantel-Cox) test was performed in GraphPad Prism testing the incremental proportion of pupae or adults per day. Supplemental Table S1 details statistical information and number of individuals analysed per replicate. Numbers were rounded to two significant figures.

## Data availability

The raw RNA sequencing data underlying Figure 2 and Supplemental Figure S2 and S3 are available in the Sequencing Read Archive (SRA) under the Bioproject ID PRJNA1155297. The R code to run statistical test is available at https://zenodo.org/doi/10.5281/zenodo.13742138.

## Acknowledgments

We would like to pay our gratitude to our colleague Jean-Géraud Issaly, who passed away in December 2022 and whose work was essential for mosquito rearing and colony maintenance in the lab. We thank Eric Marois (CNRS, INSERM, University of Strasbourg) for providing the Aaeg-M strain, and Siegfried Hapfelmeier (University of Bern) for providing the *Escherichia coli* HA 416 strain. We also thank Stencey Fontenelle and Gabrielle Georgeon (Institut Pasteur de la Guyane) for technical assistance during experiments and Emmanuel Sechet for strain handling. This study is funded by the French Government’s Investissement d’Avenir program, Laboratoire d’Excellence “Integrative Biology of Emerging Infectious Diseases” (grant ANR10-LABX-62-IBEID), and by ANR JCJC MosMi funding to M.G. (grant ANR-18-CE15-0007).

## Author Contributions

Conceptualization, Validation – O.R., J.S.S., and M.G. Methodology: O.R., J.S.S., Y.E., F.B. and M.G Software, Formal analysis– O.R. and J.S.S. Data Curation, O.R., J.S.S., Y.E. Investigation – O.R., J.S.S., C.G., Y.E., P.F.I., and M.G. Resources, F. B. Writing – Original Draft, Visualization – O.R., M.G. Writing – Review & Editing – O.R., J.S.S., C.G., F.B. and M.G. Supervision, Project administration, Funding acquisition – M.G.

## References

1. Vector-borne diseases. https://www.who.int/news-room/fact-sheets/detail/vector-borne-diseases.

2. Clements, A. N. The Biology of Mosquitoes. Volume 1. Development, Nutrition and Reproduction. vol. 1 (New York, 1992).

3. Coon, K. L., Vogel, K. J., Brown, M. R. & Strand, M. R. Mosquitoes rely on their gut microbiota for development. Mol Ecol 23, 2727 (2014).

4. Romoli, O., Schönbeck, J. C., Hapfelmeier, S. & Gendrin, M. Production of germ-free mosquitoes via transient colonisation allows stage-specific investigation of host– microbiota interactions. Nat Commun 12, (2021).

5. Correa, M. A., Matusovsky, B., Brackney, D. E. & Steven, B. Generation of axenic Aedes aegypti demonstrate live bacteria are not required for mosquito development. Nature Communications 2018 9:1 9, 1–10 (2018).

6. Wang, Y. et al. Riboflavin instability is a key factor underlying the requirement of a gut microbiota for mosquito development. Proc Natl Acad Sci U S A 118, e2101080118 (2021).

7. Serrato-Salas, J. & Gendrin, M. Involvement of Microbiota in Insect Physiology: Focus on B Vitamins. mBio 14, (2023).

8. Douglas, A. E. The B vitamin nutrition of insects: the contributions of diet, microbiome and horizontally acquired genes. Curr Opin Insect Sci 23, 65–69 (2017).

9. Dadd, R. H. Insect nutrition: current developments and metabolic implications. Annu Rev Entomol 18, 381–420 (1973).

10. Piper, M. D. Using artificial diets to understand the nutritional physiology of Drosophila melanogaster. Curr Opin Insect Sci 23, 104–111 (2017).

11. Trager, W. M. ON THE NUTRITIONAL REQUIREMENTS OF MOSQUITO LARVAE (AEDES AEGYPTI). Am J Epidemiol 22, 475–493 (1935).

12. Lea, A. O., Dimond, J. B. & Delong, D. M. A Chemically Defined Medium for Rearing Aedes aegypti Larvae. J Econ Entomol 49, 313–315 (1956).

13. Singh, K. R. P. & Brown, A. W. A. Nutritional requirements of Aedes aegypti L. J Insect Physiol 1, 199–220 (1957).

14. Lutrat, C. et al. Combining two genetic sexing strains allows sorting of non-transgenic males for Aedes genetic control. Communications Biology 2023 6:1 6, 1–12 (2023).

15. Reyda, M. R., Fugate, C. J. & Jarrett, J. T. A Complex Between Biotin Synthase and The Iron-Sulfur Cluster Assembly Chaperone HscA That Enhances In Vivo Cluster Assembly. Biochemistry 48, 10782 (2009).

16. Zhou, J. et al. Iron–sulfur biology invades tRNA modification: the case of U34 sulfuration. Nucleic Acids Res 49, 3997 (2021).

17. van Schoor, T., Kelly, E. T., Tam, N. & Attardo, G. M. Impacts of Dietary Nutritional Composition on Larval Development and Adult Body Composition in the Yellow Fever Mosquito (Aedes aegypti). Insects 11, 1–15 (2020).

18. Chambers, G. M. & Klowden, M. J. CORRELATION OF NUTRITIONAL RESERVES WITH A CRITICAL WEIGHT FOR PUPATION IN LARVAL AEDES AEGYPTI iVIOSqUITONS.

19. Vendl, T., Šípek, P., Kouklík, O. & Kratochvíl, L. Hidden complexity in the ontogeny of sexual size dimorphism in male-larger beetles. Scientific Reports 2018 8:1 8, 1–10 (2018).

20. Seong, K. H. & Kang, S. Noncanonical function of the Sex lethal gene controls the protogyny phenotype in Drosophila melanogaster. Scientific Reports 2022 12:1 12, 1–9 (2022).

21. Souza, R. S. et al. Microorganism-based larval diets affect mosquito development, size and nutritional reserves in the yellow fever mosquito aedes aegypti (Diptera: Culicidae). Front Physiol 10, (2019).

22. Yan, J., Kim, C. H., Chesser, L., Ramirez, J. L. & Stone, C. M. Nutritional stress compromises mosquito fitness and antiviral immunity, while enhancing dengue virus infection susceptibility. Communications Biology 2023 6:1 6, 1–15 (2023).

23. Matthews, B. J. et al. Improved reference genome of Aedes aegypti informs arbovirus vector control. Nature 563, 501–507 (2018).

24. Goos, J. M., Cameron, |, Swain, J., Munch, S. B. & Walsh, M. R. Maternal diet and age alter direct and indirect relationships between life-history traits across multiple generations. (2018) doi:10.1111/1365-2435.13258.

25. Deas, J. B., Blondel, L. & Extavour, C. G. Ancestral and offspring nutrition interact to affect life-history traits in Drosophila melanogaster. Proceedings of the Royal Society B: Biological Sciences 286, (2019).

26. Neophytou, C. & Pitsouli, C. Biotin controls intestinal stem cell mitosis and host-microbiome interactions. Cell Rep 38, (2022).

27. Velazquez-Arellano, A. & Hernandez-Vazquez, A. D. J. Vitamins as Cofactors for Energy Homeostasis and Their Genomic Control, With Special Reference to Biotin, Thiamine, and Pantothenic Acid. Principles of Nutrigenetics and Nutrigenomics: Fundamentals of Individualized Nutrition 271–277 (2020) doi:10.1016/B978-0-12-804572-5.00035-5.

28. Trager, W. Biotin And Fat-Soluble Materials With Biotin Activity In The Nutrition Of Mosquito Larvae. Journal of Biological Chemistry 176, 1211–1223 (1948).

29. Pillai, M. K. K. & Madhukar, B. V. R. Effect of biotin on the fertility of the yellow fever mosquito, Aedes aegypti (L.). Naturwissenschaften 56, 218–219 (1969).

30. Fraenkel, G. & Blewett, M. The vitamin B-complex requirements of several insects. Biochemical Journal 37, 686 (1943).

31. Benschoter, C. A. & Paniagua G., R. Reproduction and Longevity of Mexican Fruit Flies, Anastrepha ludens (Diptera: Tephritidae), Fed Biotin in the Diet. Ann Entomol Soc Am 59, 298–300 (1966).

32. Benschoter, C. A. Effect of Dietary Biotin on Reproduction of the House Fly. J Econ Entomol 60, 1326–1328 (1967).

33. Cohen, E. & Levinson, H. Z. Disrupted fertility of the hidebeetle Dermestes maculatus (Deg.) due to dietary overdosage of biotin. Experientia 24, 367–368 (1968).

34. Levinson, H. Z. & Cohen, E. The action of overdosed biotin on reproduction of the hide beetle, Dermestes maculatus. J Insect Physiol 19, 551–558 (1973).

35. KEGG PATHWAY: Biotin metabolism - Microbacterium sp. SGAir0570. https://www.kegg.jp/pathway/mics00780.

36. Scholz, M. et al. Large scale genome reconstructions illuminate Wolbachia evolution. Nat Commun 11, (2020).

