## Supplemental for "Nutritional sex-specificity on bacterial metabolites during mosquito development leads to adult sex-ratio distortion"

### Supplemental results

*Transcriptomic analysis of germ-free larvae treated with B vitamins*

We conducted a transcriptomic study on larvae cleared of their microbiota at the beginning of the third instar and kept in germ-free conditions (GF), supplemented with VIT1x or VIT8x solutions. When transferring third instar larvae in a medium without *m*DAP and D-Ala to derive them germ-free, we supplemented their diet with 0x, 1x or 8x concentrations of B vitamins (Supplemental Figure S2A). As the mortality was generally observed at the end of the fourth instar or during metamorphosis, we decided to sample individuals earlier, 3-6 hours after moulting to the fourth instar. We extracted RNA and DNA from individual larvae (1200 larvae in total) and sexed them by individual PCR on the male-specific *Nix* gene. The RNA of at least 48 females and males per replicate was pooled in equimolar amount. The large number of samples imposed experimental set up adjustments which resulted in more than one freeze-thaw cycle for each sample, hence the resulting RNA quality did not allow to proceed for poly-A enrichment for RNA sequencing. Instead, libraries were prepared by depleting ribosomal RNA, which allowed us to sequence non-coding RNAs together with protein-coding transcripts. After quality trimming, reads were aligned on the *Ae. aegypti* genome. An average of 45 million reads was obtained per sample (min 22 million, max 70 million), with an average alignment rate of 80 % (min 6 %, max 90 %). Ribosomal RNA reads composed less than the 0.4 % of all reads, indicating a successful depletion of these RNA species.

As a quality control for the sex assignment of our samples, we quantified the amount of reads mapping on the *Nix* gene in each sample (AAEL0922912, Supplemental Figure S2B). Read counts were always lower in females than in males, except for the third replicate of the germ-free sample. A second check on our PCR results spotted several mistakes in this replicate, thus all samples belonging to the third replicate were removed from the analysis (Supplemental Figure S2A). Moreover, the sample of males treated with VIT1x collected during the first replicate showed very low alignment rates (5.6 %) and only one read aligning on the *Nix* gene, making any analyses impossible on the VIT1x condition. In summary, we performed the differential expression analysis only on germ-free and VIT8x treatments using the first and second replicates. The general low RNA quality and the suboptimal number of replicates resulted in an overall poor clustering of samples based on sex or treatment (Supplemental Figure S2C). Due to a low number of regulated genes, we could not perform any Gene Ontology enrichment analysis.

To verify the accuracy of sample sex assignment, we first compared the transcriptomes of male and female larvae. Principal component analysis (PCA) discriminated male and female samples in germ-free treated-larvae but not in VIT8x conditions (Supplemental Figure S2D, E), and discriminated germ-free and VIT8x conditions in females but not in males (Supplemental Figure S2F, G). A total number of 69 and 20 genes were differentially expressed between males and females in germ-free or VIT8x conditions, respectively (Supplemental Figure S3A, Supplemental Tables S2 and S3). When comparing which differentially regulated genes were commonly identified in both germ-free and VIT8x conditions, five and three genes were up- and down-regulated in males compared to females. Among these, the male determiner gene *Nix* (Hall et al. 2015) was significantly up-regulated in males in both GF and VIT8x samples, and the male-specific gene *myo-sex* (Aryan et al. 2020) was among the up-regulated genes in males kept in GF conditions, corroborating the sex-specificity of the genes identified as such in our data.

While PCA did not clearly separate samples for the vitamin treatment (Supplemental Figure S4D, E), a small number of genes were identified as differentially regulated by the vitamin treatment, with 13 and 16 genes significantly regulated in males and females, respectively (Supplemental Figure S3C, D). The *AAEL020376* gene, paralog to the *chloride channel protein 2* gene, was commonly down-regulated with vitamins in both sexes, suggesting its involvement in a generalist vitamin-regulated mechanism (Supplemental Figure S3C, D). Amongst the genes induced in males upon treatment, heat shock protein 70 encoding gene was highly up-regulated, indicating a possible activation of a stress-response in vitamin-treated male larvae (Figure 2B, *padj* < 0.001). Most of these genes were not annotated in the *Ae. aegypti* genome, but some had known orthologs. *AAEL004564* is an ortholog of *CG8420* in *Drosophila*, which encodes a male accessory gland protein, putatively component of the seminal fluid (Ram and Wolfner 2007). *AAEL013606* (Figure 2C, *padj* = 0.004) is a sex-determining region Y protein (SRY)-like ortholog, a DNA-binding transcription factor that initiates male sex-determination in most mammals. Others appear to have more tangential male-specific functions, such as *AAEL002537*, ortholog to the *Drosophila* gene *dusky*, which is involved in cell morphogenesis and imaginal disc-derived wing morphogenesis. *AAEL022708* is annotated as *Peptidyl-prolyl cis-trans isomerase*, a chaperone protein that regulates protein conformational changes by catalysing the isomerization between the *cis* and *trans* forms of peptide bonds. Its *Drosophila* ortholog (*CG11858*) is highly expressed in the larval central nervous system and is predicted to bind DNA and process rRNA.

Female larvae treated with vitamins showed a higher number of down-regulated genes, including a lipase 1 precursor encoding gene (Figure 2D, *padj* = 0.04) and two genes involved in ion transport across membranes (*AAEL010140* and *AAEL011109*). Among the up-regulated genes, three were orthologs to genes encoding for chitin binding proteins expressed during development (*AAEL024503*, *AAEL006686* and *AAEL017212*, Figure 2E, *padj* = 0.02). Even though these results do not allow to draw strong conclusions, the upregulation in structural genes and down regulation in lipase may reflect a positive impact on developmental processes, while the downregulation of ion transport may be a response to stress due to increasing concentrations of vitamins.

### Supplemental Materials and Methods

*Experimental setup and sample collection for transcriptomics on germ-free vs vitamin treated larvae*

Germ-free third instar larvae were obtained as described above. To maximise the number of larvae, at day four (day five for the Aaeg-M strain) plates were inspected every 3 h and newly moulted third instar larvae were washed and transferred in new 24-well plates filled either with sterile water (germ-free condition), or with a 1x or 8x B vitamin solution (i.e., a solution of B vitamins each at the reference concentration or at 8 times the reference concentration). The following day (day five or day six for the Aaeg-M strain), wells of fourth instar larvae were marked every 3 h and the corresponding larvae were sampled 3 h later. This experimental set up was chosen to sample "synchronized" larvae that are all in the same development timing (3 to 6 h after their moulting to fourth instar) even though they have been sampled over a 12 h period. Individual larvae were placed in individual screwcap 2 mL tubes filled with glass beads and immediately stored at -80 °C. Three independent replicates were performed. For each replicate, 48 random larvae per condition were left in the plates (i.e., not sampled) and observed for 14 days to analyse their developmental success. Up to 150 larvae per condition and replicate were collected for RNA extraction and transcriptome analysis.

*DNA and RNA extraction of transcriptomic samples*

DNA and RNA were simultaneously extracted from individual larvae using Trizol (Thermo Fisher Scientific). Individual larvae were homogenised in 500 µL of Trizol using a bead beater (Precellys Evolution, Bertin) for 2x1 min at 9,000 rpm. After adding chloroform and centrifuging, RNA was extracted from the upper aqueous phase following manufacturer's instructions, resuspended in 20 µL of nuclease free water and stored at -80 °C. In parallel, DNA was purified from the bottom organic phase by precipitation using 0.1 M sodium citrate and following manufacturer's instructions. DNA was resuspended in 8 mM NaOH and stored at -20 °C until use.

*Determination of sex of individual larvae by PCR for transcriptomics analysis*

Larvae were sexed by multiplex PCR on *Nix* (present only in males) and *actin* (reaction positive control). The primers used are listed in Supplementary Table 4. Each PCR was set-up with 1 U of FirePol DNA Polymerase (Solis Biodyne), 1x Buffer BD, 0.2 mM dNTPs, 500 nM of each primer, 1.5 mM $MgCl_2$ and 2 µL of DNA. The amplification cycle included an initial denaturation at 95 °C for 5 min, followed by 35 cycles at 95 °C for 20 sec, at 59 °C for 40 sec and 72 °C for 30 sec and a final 5 min elongation step at 72 °C. PCR products were visualised using a QIAxcel DNA screening kit in a QIAxcel instrument (Qiagen).

*Pooling of RNA and sequencing*

After determination of larval sex by PCR on DNA, RNA was pooled for males and females independently. Pooled samples were treated for 30 min at 37 °C with 10 U of DNase I (Thermo Fisher Scientific) to remove any DNA trace and purified using the MagJET RNA Kit (Thermo Fisher Scientific) in a KingFisher Duo Prime system (Thermo Fisher Scientific). Samples were mixed with RNA stable (Biomatrica), vacuum dried using a SpeedVac and shipped for library preparation and sequencing.

*Analysis of RNA sequencing*

RNA quality evaluation, library preparation and sequencing were performed by Microsynth (Balgach, Switzerland). RNA integrity was evaluated on a BioAnalyzer (Agilent Technologies) using an RNA 6000 Pico RNA chip. Paired-end libraries (2x50 bp) were constructed using the Universal RNA-Seq with NuQuant kit (Nugen Tecan Genomics) from 250 or 45 ng of total RNA. *Ae. aegypti* ribosomal RNA was depleted using a custom AnyDeplete probe mix (Nugen Tecan Genomics). After quality check, libraries were pooled in equimolar concentrations and sequenced on a NovaSeq SP flow cell (Illumina). The resulting raw reads were de-multiplexed and adaptor sequences were trimmed by the sequencing facility. FASTQ sequences were trimmed for their quality using Trimmomatic (Bolger et al. 2014)(minimum quality score: 15, minimal length: 40) and their quality was evaluated using FASTQC(Babraham Bioinformatics). After quality trimming, an average of 45 million reads (min 22 million, max 70 million) were obtained per sample. Reads were mapped on the *Ae. aegypti* genome (assembly AeaegL5, geneset AeaegL5.3) using HISAT2 (Kim et al. 2019). The average percentage of reads aligned concordantly one time was 49% (min 3.6%, max 58%). Read counts were obtained with FeatureCounts (Liao et al. 2014)and differential expression analysis was conducted in R using the DESeq2 package (Love et al. 2014).

### Supplemental Figures

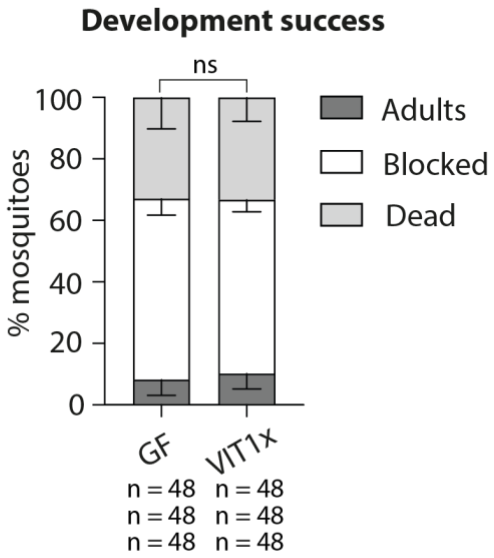

**Supplemental Figure S1. Development success of mosquitoes exposed to B vitamins at 1x concentrations during larval development.** Proportion of mosquitoes completing their development (dark grey), blocked at the larval stage (white), or dying (light grey) when reversibly colonized until the beginning of the third instar and kept in standard germ-free conditions (GF) or supplemented with a 1x (VIT1x) B vitamin solution. Bar charts represent the mean ± SEM of three independent replicates. Numbers below graphs indicate the number of mosquitoes analysed per replicate and condition.

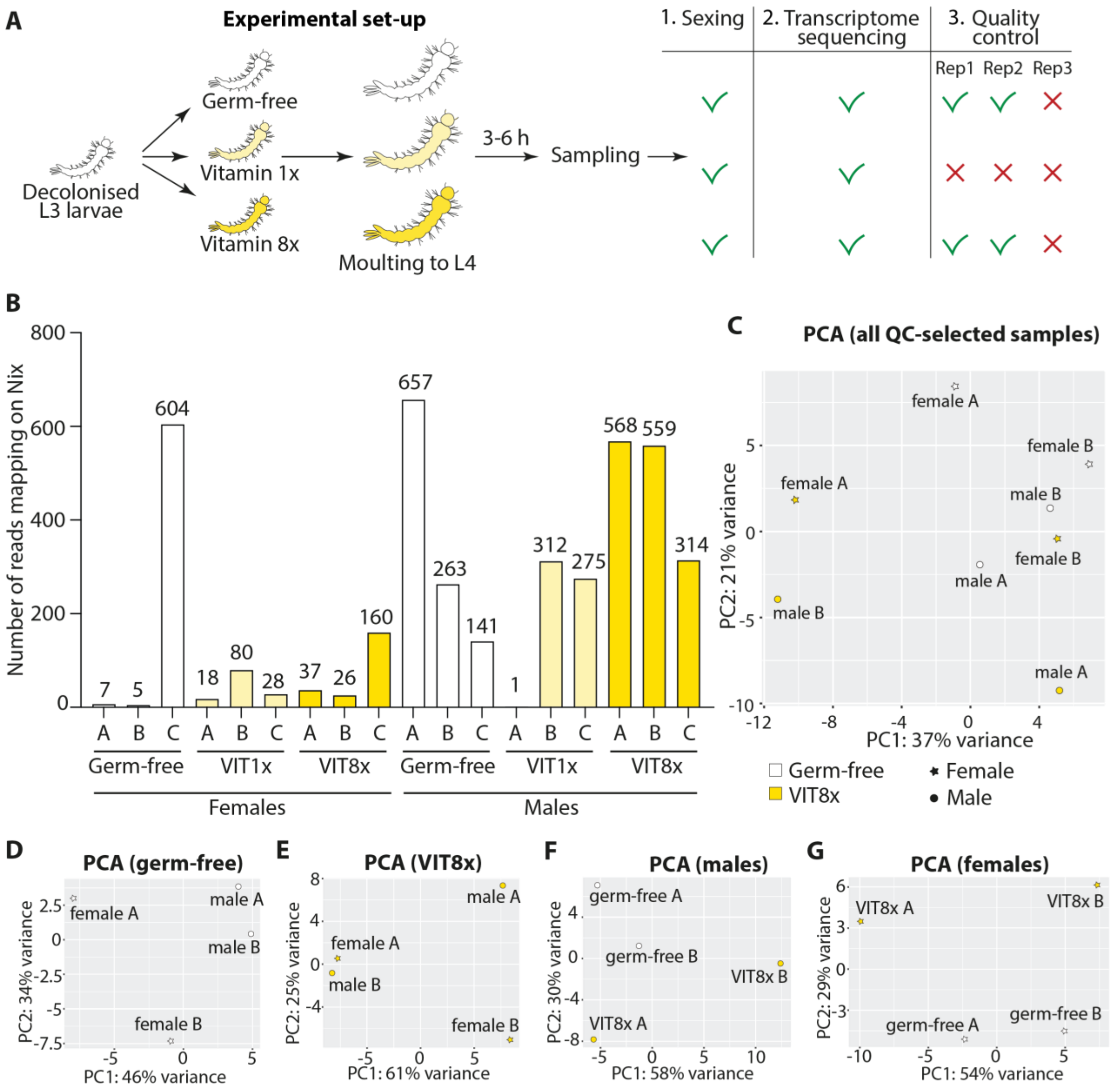

**Supplemental Figure S2. Experimental set-up and quality check of transcriptomic analysis of male and female larvae exposed to B vitamins.** (**A**) Experimental set-up: mosquito larvae reared in gnotobiotic conditions with the auxotrophic *E. coli* strain were decolonized at the beginning of the third instar (L3) to obtain germ-free larvae. Larvae were either kept in germ-free conditions or treated with 1x or 8x B vitamin solutions. After three to six hours since moulting to the fourth instar (L4), up to 150 larvae per condition were individually collected. Three independent replicates were performed. DNA was extracted from individual larvae for sex assignment using the *Nix* gene, while RNA was extracted for transcriptome sequencing on male and female larvae. A quality control step on the proportion of *Nix* reads and of overall alignment rates and a second check of initial *Nix* PCR results, allowed to select samples for data analysis. (**B**) Raw counts of read mapping on the *Nix* gene in the sequenced transcriptomes. Total number of reads mapping on the *Nix* gene in the transcriptomes of female and male larvae reversibly colonized at the beginning of the third instar and kept in germ-free conditions (GF) or supplemented with a 1x (VIT1x) or 8x (VIT8x) B vitamin solution. Results from the three replicates (A, B, and C) are shown separately. (**C-G**) Principal Component Analysis (PCA) of transcriptomes passing the quality check and used for the final analysis. (**C**) PCA of transcriptomes of female (star) and male (circle) larvae kept in germ-free conditions (white) or treated with vitamins (VIT8x, yellow). (**D, E**) PCA of transcriptomes of female (star) and male (circle) larvae kept in germ-free conditions (**D**) or treated with vitamins (**E**). (**F, G**) PCA of transcriptomes of male (**F**) and female (**G**) larvae kept in germ-free conditions (white) or treated with vitamins (yellow).

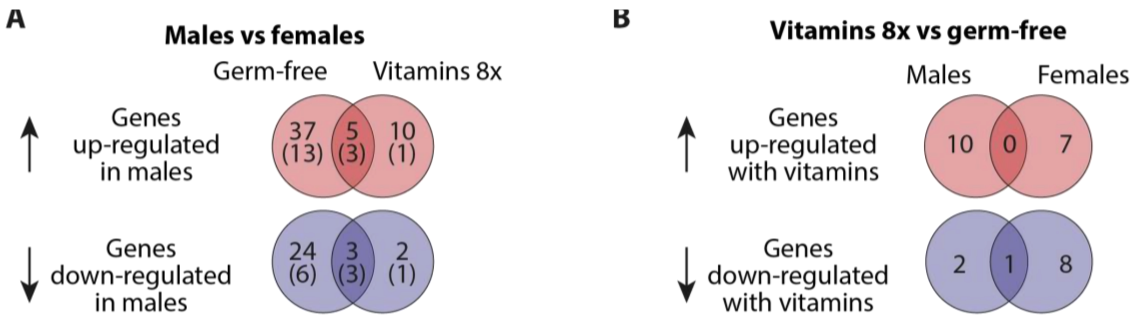

**C** **Males with vitamins 8x - Differentially regulated genes**

| ID | log2FoldChange | padj | Transcript product | Transcript type | Paralog/Ortholog, species (description) |
|---|---|---|---|---|---|
| AAEL021701 | -2.72 | 2.8E-03 | unspecified product | Coding gene | cuticle protein 63-like, *Aedes albopictus* |
| **AAEL020376** | **-2.43** | **2.2E-03** | **unspecified product** | Coding gene | **chloride channel protein 2, *Aedes aegypti*** |
| AAEL028740 | -1.96 | 6.8E-08 | unspecified product | ncRNA | |
| AAEL007377 | 1.01 | 1.9E-02 | ammonium transporter | Coding gene | ammonium transporter, *Aedes aegypti* |
| AAEL010390 | 1.27 | 3.0E-02 | glucosyl/glucuronosyl transferases | Coding gene | glucosyl/glucuronosyl transferase, *Aedes aegypti* |
| AAEL022253 | 1.44 | 1.3E-02 | unspecified product | Pseudogene | |
| AAEL017976 | 1.51 | 3.4E-05 | heat shock protein HSP70 | Coding gene | heat shock protein HSP70, *Aedes aegypti* |
| AAEL004564 | 2.02 | 5.1E-03 | unspecified product | Coding gene | CG8420, *Drosophila melanogaster* (involved in sexual reproduction*) |
| AAEL013606 | 2.85 | 3.8E-03 | unspecified product | Coding gene | sex-determining region Y (SRY) protein-like, *Aedes albopictus* |
| AAEL002537 | 2.88 | 9.1E-03 | unspecified product | Coding gene | dusky, *Drosophila melanogaster* (regulates epidermal cell size/shape changes during wing morphogenesis*) |
| AAEL027398 | 3.82 | 5.3E-03 | unspecified product | Coding gene | DUF4773 domain-containing protein, *Anopheles albimanus* |
| AAEL000484 | 4.71 | 2.8E-03 | unspecified product | Coding gene | prespore vesicle protein-like, *Aedes albopictus* |
| AAEL022708 | 6.76 | 1.9E-02 | Peptidyl-prolyl cis-trans isomerase | Coding gene | CG11858, *Drosophila melanogaster* (predicted to enable DNA binding activity, involved in rRNA processing*) |

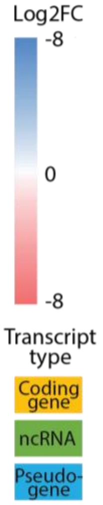

**D** **Females with vitamins 8x - Differentially regulated genes**

| ID | log2FoldChange | padj | Transcript product | Transcript type | Paralog/Ortholog, species (description) |
|---|---|---|---|---|---|
| AAEL012340 | -8.01 | 3.8E-02 | lipase 1 precursor | Coding gene | lysosomal acid lipase, putative, *Aedes aegypti* |
| **AAEL020376** | **-2.93** | **6.8E-03** | **unspecified product** | Coding gene | **chloride channel protein 2, *Aedes aegypti*** |
| AAEL017678 | -2.47 | 1.9E-02 | U4 spliceosomal RNA | ncRNA | |
| AAEL010140 | -1.71 | 3.9E-02 | sodium/potassium-dependent ATPase beta-2 subunit | Coding gene | sodium/potassium-transporting ATPase subunit beta-1-like, *Aedes albopictus* |
| AAEL017646 | -1.53 | 1.1E-03 | U1 spliceosomal RNA | ncRNA | |
| AAEL017691 | -1.21 | 3.9E-02 | U1 spliceosomal RNA | ncRNA | |
| AAEL011109 | -1.13 | 3.8E-02 | unspecified product | Coding gene | sodium/hydrogen exchanger 9B2, *Aedes albopictus* |
| AAEL028652 | -1.10 | 1.4E-02 | unspecified product | ncRNA | |
| AAEL017826 | -0.96 | 3.9E-02 | U1 spliceosomal RNA | ncRNA | |
| AAEL017212 | 0.99 | 3.6E-02 | unspecified product | Coding gene | acidic leucine-rich nuclear phosphoprotein 32 family member B-like, *Aedes albopictus* |
| AAEL028790 | 1.17 | 3.8E-02 | unspecified product | ncRNA | |
| AAEL017067 | 1.34 | 1.6E-02 | unspecified product | Coding gene | peritrophin-48-like, *Aedes albopictus* |
| AAEL006686 | 1.49 | 3.8E-02 | unspecified product | Coding gene | CG43294, *Drosophila melanogaster* (predicted to enable chitin binding activity*) |
| AAEL024503 | 1.68 | 9.2E-07 | unspecified product | Coding gene | AAEL024503, *Aedes albopictus* (contains a chitin binding Peritrophin-A domain) |
| AAEL028730 | 3.30 | 9.1E-24 | unspecified product | ncRNA | |
| AAEL022711 | 6.51 | 1.6E-02 | unspecified product | ncRNA | |

**Supplemental Figure S3. Genes differentially regulated in male and female larvae exposed to B vitamins.** (**A**) Number of genes up- (red) and downregulated (blue) in male vs female mosquitoes when larvae were kept in germ-free conditions or treated with the 8x B vitamin solution. The numbers in parenthesis represent the number of genes that are shared between our study and the Matthews *et al* transcriptome (Matthews et al. 2018). (**B**) Number of genes up- (red) and downregulated (blue) in B vitamin vs germ-free conditions in male and female larvae. (**C, D**) List of genes up- (red) and down-regulated (blue) in male (**C**) and female (**D**) larvae treated with the B vitamin solution. For each gene, Vectorbase ID, $log_2$ fold value, *padj*, description, and transcript type (coding gene: yellow; noncoding RNA (ncRNA): green; pseudogene: blue) are indicated. The last column shows paralog or ortholog genes with the name of the species and a brief function description. Asterisks (*) indicate description copied from FlyBase.

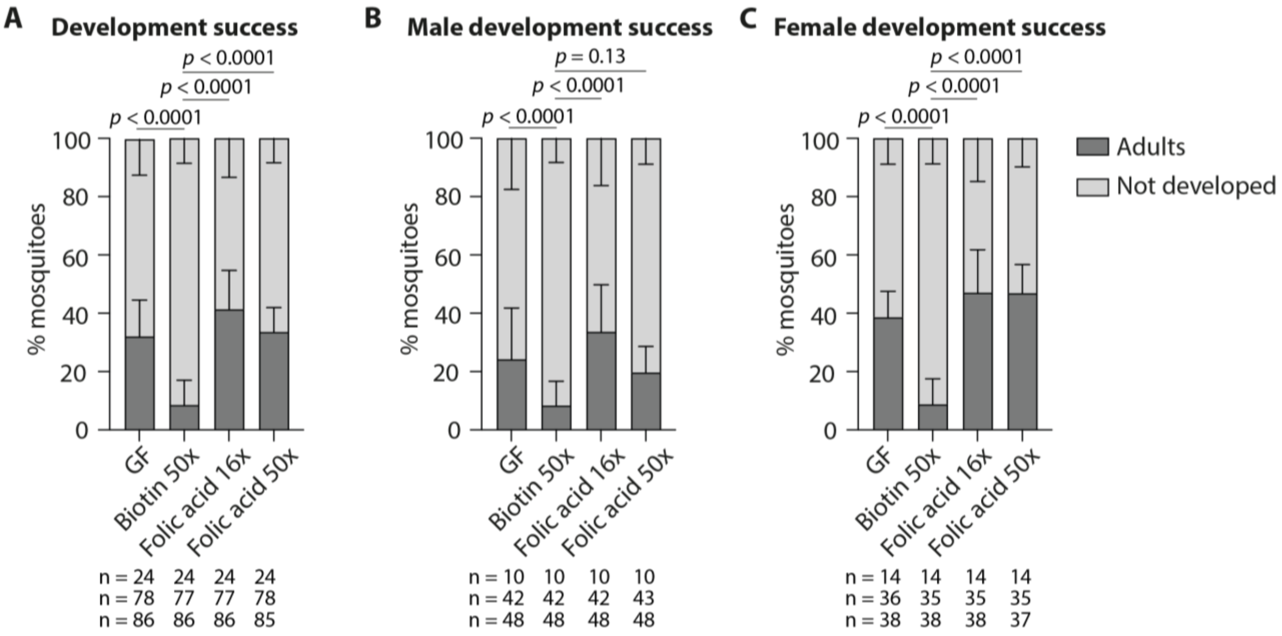

**Supplemental Figure S4. Development success of male and female mosquito larvae exposed to high doses of biotin and folic acid separately.** Proportion of mosquitoes (Aaeg-M strain) completing their development (dark grey), or not developing (light grey) when reversibly colonized at the beginning of the third instar and kept in germ-free conditions (GF) or provided with a 50x biotin solution, or a 16x or 50x folic acid solution. Bar charts represent the mean ± SEM of three independent replicates. Data in (**A**) are not sorted by sex, while data in (**B**) and (**C**) represent results for male and female mosquitoes, respectively. Numbers below graphs indicate the number of mosquitoes analysed per replicate and condition. See Supplemental Table S1 for detailed statistical information.

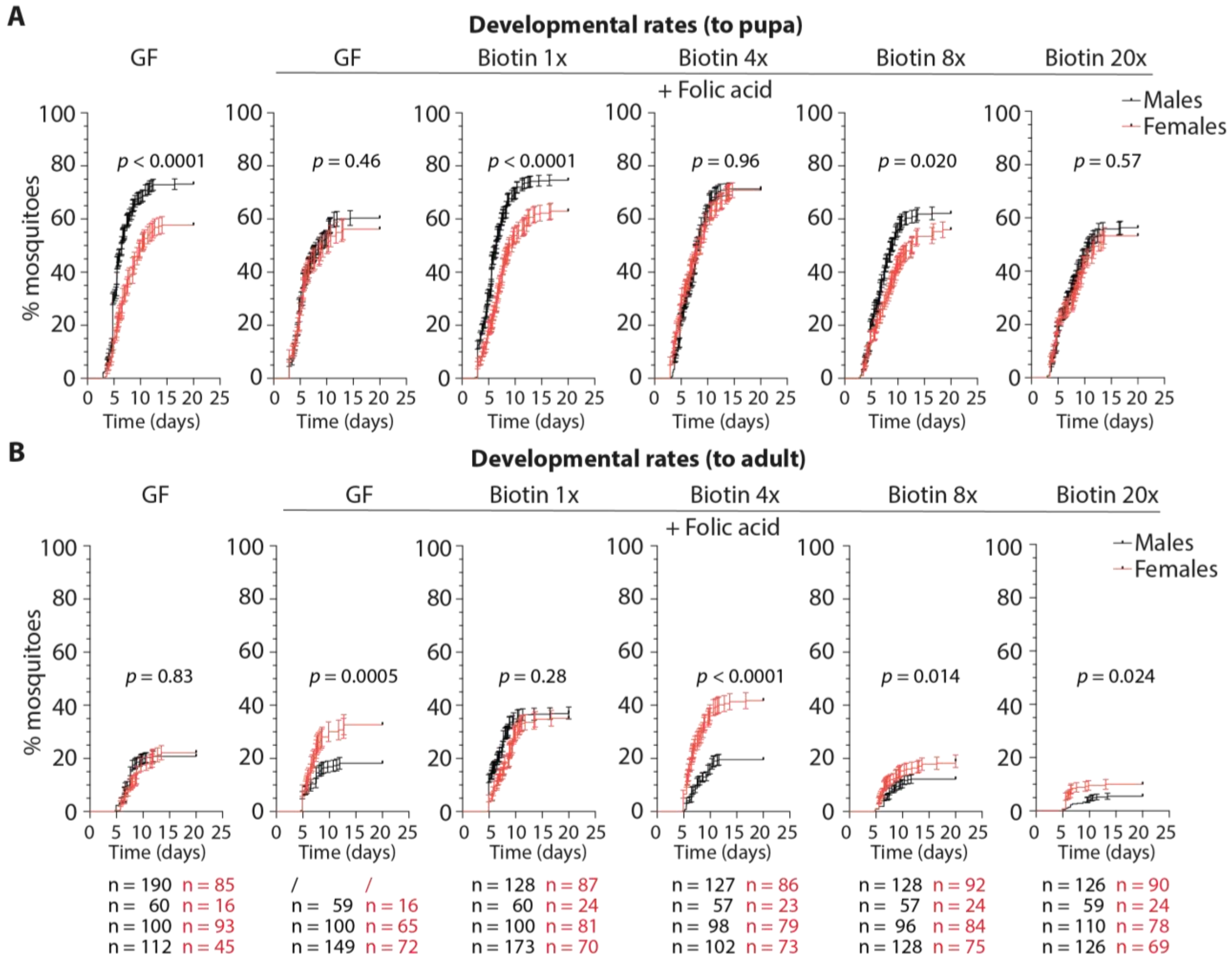

**Supplemental Figure S5. Developmental rates to pupa or adult of mosquitoes exposed to increasing doses of biotin during larval development.** Proportion of male (black) and female (red) larvae becoming pupae (**A**) or adult (**B**) per day when reversibly colonized at the beginning of the third instar and kept in germ-free conditions (GF) or provided with a 1x, 4x, 8x, or 20x biotin solution. A 16x folic acid solution was also added to all tested conditions. Each dot represents the mean ± SEM of four independent replicates except for GF+folic acid (three replicates). Numbers below graphs indicate the number of mosquitoes analysed per replicate and condition. See Supplementary Table 1 for detailed statistical information.

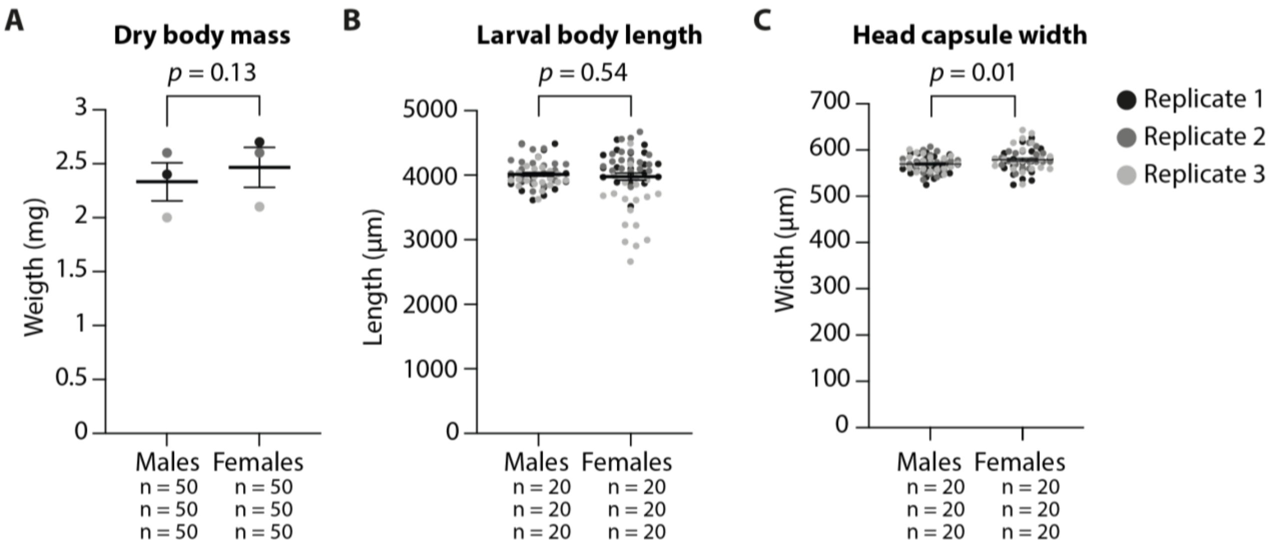

**Supplemental Figure S6. Differences in body weight and size between male and female larvae.** Measurements were conducted on third instar larvae (Aaeg-M strain) reared in gnotobiotic conditions. (**A**) Comparison of the dry body mass of pools of 50 male and female larvae. (**B-C**) Estimates of male and female larvae size as body length (**B**) and head capsule width (**C**). Three replicates were performed for each measurement. Numbers below graphs indicate the number of mosquitoes analysed per replicate and sex.

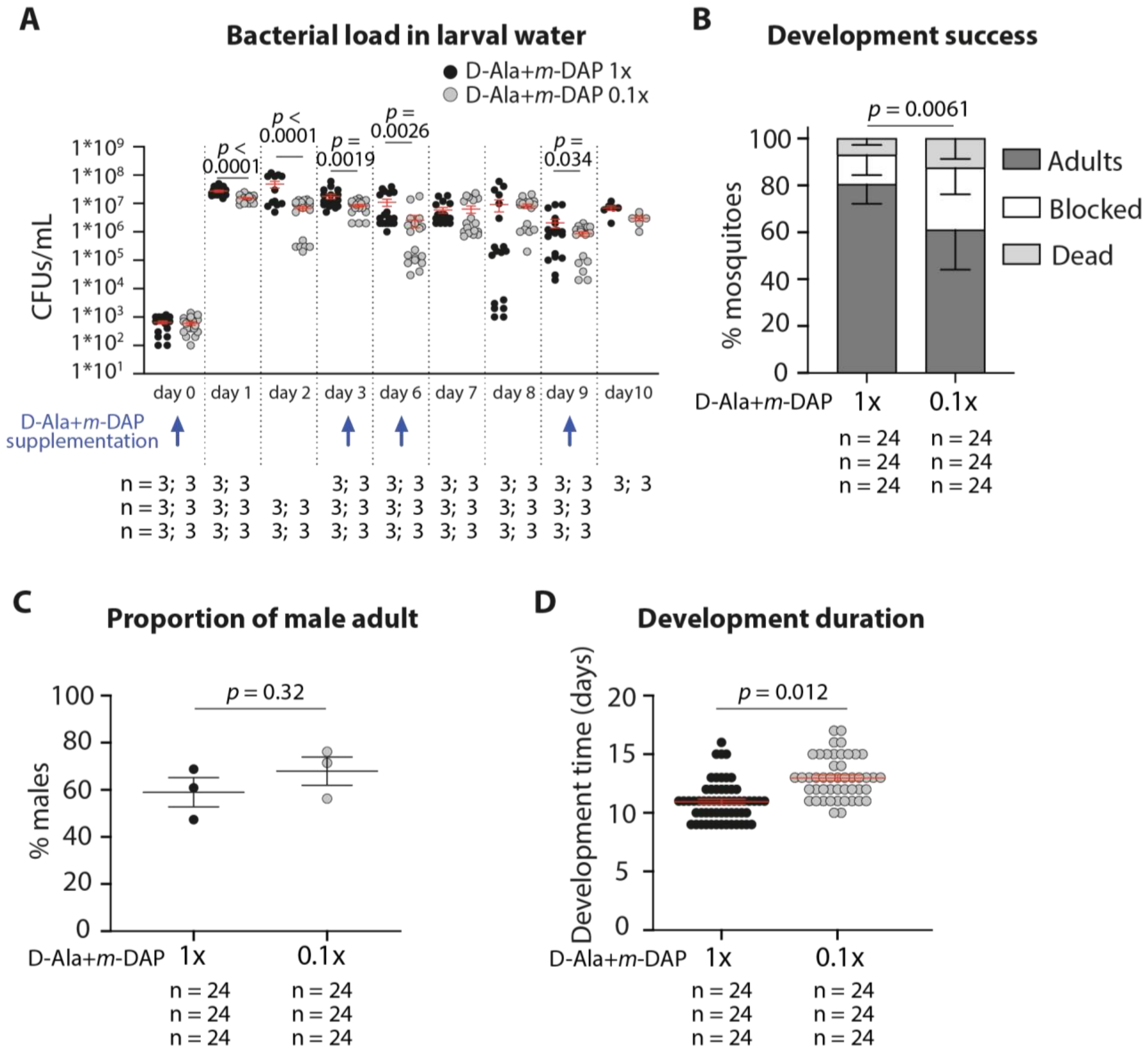

**Supplemental Figure S7. Development success and sex ratio of larvae with decreased number of bacteria.** In **A**-**D**, auxotrophic *E. coli*-colonized larvae were treated with 1x or 0.1x concentrations of D-Ala and m-DAP. (**A**) Number of CFUs/mL in wells containing individual larvae. Each dot represents a well, 6 wells/time point/replicate were tested in each replicate. Time-points indicate the time after bacteria were added to germ-free larvae. (**B**) Proportion of mosquitoes reaching adulthood (dark grey), blocked in development (white) or dead (light grey) when auxotrophic *E. coli*-colonized larvae were supplemented with standard (1x) or diluted (0.1x) D-Ala and *m*-DAP concentrations. (**C**) Proportion of males amongst the adults and (**D**) number of days until reaching adulthood when auxotrophic *E. coli*-colonized larvae were supplemented with standard (1x) or diluted (0.1x) D-Ala and *m*-DAP concentrations.

Romoli_Serrato-Salas_SuppFig8

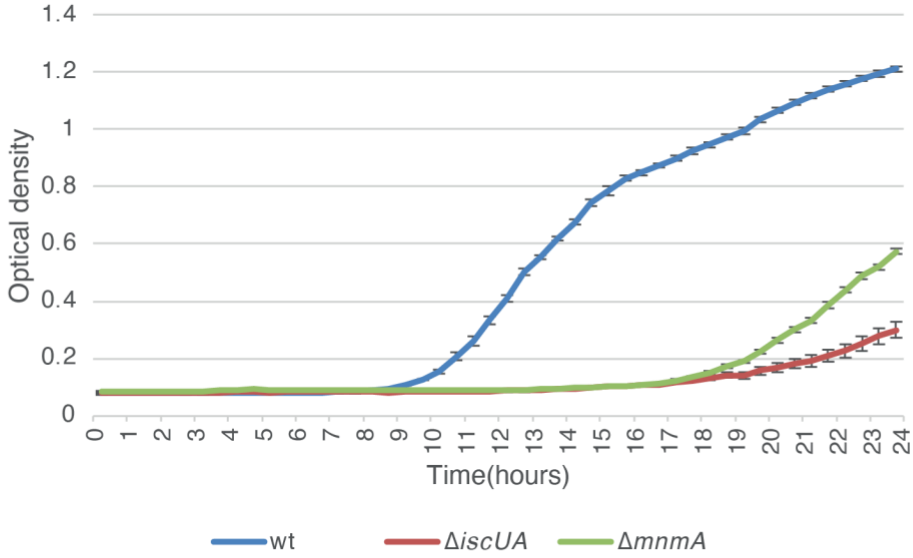

**Figure S8. Growth curves of wt *E. coli*, growth deficient mutant Δ*mnmA* or biotin-deficient mutant Δ*iscUA*.** Optical density of individual wells of a 96-well plate, measured every 30 min during 23 h 30 min. For each bacterium, 3 independent colonies were used to start the pre-cultures, and pre-cultures were diluted to 1:50 as duplicates to start the experiment. Data show the average ± SEM of the six wells per strain per time point.

### Supplementary Table S1. Detailed results of statistical tests.

| Analysis | Response variable | Predictor | Random effect | Sample Size | Test result | Comparisons | | |
|---|---|---|---|---|---|---|---|---|
| **Figure 1A** | | | | | | | | |
| Glmm (binomial) | % adults | Treatment | Replicate | **Replicate A:** AUX (n=48); GF (n=57); VIT4a (n=48); VIT8a (n=48) **Replicate B:** AUX (n=48); GF (n=48); VIT4a (n=48); VIT8a (n=48) **Replicate C:** AUX (n=47); GF (n=24); VIT4a (n=24); VIT8a (n=48) | $F_3 = 24.11$, $p = 0.0001$ | lsmeans (Bonferroni correction):<br>AUX vs GF: $p = 0.0001$<br>AUX vs VIT4a: $p = 0.0001$<br>AUX vs VIT8a: $p = 0.0001$<br>GF vs VIT4a: $p = 0.71$<br>GF vs VIT8a: $p = 0.58$<br>VIT4a vs VIT8a: $p = 0.0057$ | | |
| | % dead | Treatment | Replicate | | $F_3 = 22.01$, $p < 0.0001$ | lsmeans (Bonferroni correction):<br>AUX vs GF: $p < 0.0001$<br>AUX vs VIT4a: $p < 0.0001$<br>AUX vs VIT8a: $p < 0.0001$<br>GF vs VIT4a: $p = 0.012$<br>GF vs VIT8a: $p = 0.0001$<br>VIT4a vs VIT8a: $p = 0.67$ | | |
| | % blocked | Treatment | Replicate | | $F_3 = 18.34$, $p = 0.0001$ | lsmeans (Bonferroni correction):<br>AUX vs GF: $p = 1.0$<br>AUX vs VIT4a: $p = 1.0$<br>AUX vs VIT8a: $p = 1.0$<br>GF vs VIT4a: $p < 0.0001$<br>GF vs VIT8a: $p = 0.0013$<br>VIT4a vs VIT8a: $p = 0.048$ | | |
| **Figure 1B** | | | | | | | | |
| Glmm (binomial) | Sex | Treatment | Replicate | **Replicate A:** AUX (n=48); GF (n=47); VIT4a (n=48); VIT8a (n=48) **Replicate B:** AUX (n=48); GF (n=48); VIT4a (n=48); VIT8a (n=48) **Replicate C:** AUX (n=47); GF (n=24); VIT4a (n=24); VIT8a (n=48) | $F_3 = 8.08$, $p < 0.0001$ | lsmeans (Bonferroni correction):<br>AUX vs GF: $p = 1.0$<br>AUX vs VIT4a: $p = 0.0008$<br>AUX vs VIT8a: $p = 0.0037$<br>GF vs VIT4a: $p = 0.0005$<br>GF vs VIT8a: $p = 0.0019$<br>VIT4a vs VIT8a: $p = 1.0$ | | |
| **Figure 3A** | | | | | | | | |
| Glmm (binomial) | % adults | Treatment | Replicate | **Replicate A:** GF (n=24); Folic acid (n=24); Choline (n=24); Thiamine (n=24); Nicotinic acid (n=24); Biotin (n=24); Pyridoxine (n=24) **Replicate B:** GF (n=24); Folic acid (n=24); Choline (n=24); Thiamine (n=24); Nicotinic acid (n=24); Biotin (n=24); Riboflavin (n=24); Pyridoxine (n=24) **Replicate C:** GF (n=24); Folic acid (n=24); Choline (n=24); Thiamine (n=24); Nicotinic acid (n=24); Biotin (n=24); Riboflavin (n=24); Pyridoxine (n=24) | $F_7 = 12.30$, $p = 0.0001$ | lsmeans (Bonferroni correction):<br>GF vs Biotin: $p = 0.0001$<br>GF vs Choline: $p = 1.0$<br>GF vs Folic acid: $p = 0.015$<br>GF vs Nicotinic acid: $p = 1.0$<br>GF vs Pyridoxine: $p = 0.58$<br>GF vs Riboflavin: $p = 0.21$<br>GF vs Thiamine: $p = 1.0$<br>Biotin vs Choline: $p = 0.0001$ | Biotin vs Folic acid: $p = 0.0001$<br>Biotin vs Nicotinic acid: $p = 0.0001$<br>Biotin vs Pyridoxine: $p = 0.0001$<br>Biotin vs Riboflavin: $p < 0.0001$<br>Biotin vs Thiamine: $p < 0.0001$<br>Choline vs Folic acid: $p = 0.062$<br>Choline vs Nicotinic acid: $p = 1.0$<br>Choline vs Pyridoxine: $p = 1.0$<br>Choline vs Riboflavin: $p = 0.58$<br>Choline vs Thiamine: $p = 1.0$ | Folic acid vs Nicotinic acid: $p = 0.12$<br>Folic acid vs Pyridoxine: $p = 1.0$<br>Folic acid vs Riboflavin: $p = 1.0$<br>Folic acid vs Thiamine: $p = 1.0$<br>Nicotinic acid vs Pyridoxine: $p = 1.0$<br>Nicotinic acid vs Riboflavin: $p = 0.91$<br>Nicotinic acid vs Thiamine: $p = 1.0$<br>Pyridoxine acid vs Riboflavin: $p = 1.0$<br>Pyridoxine acid vs Thiamine: $p = 1.0$<br>Riboflavin acid vs Thiamine: $p = 1.0$ |
| | % dead | Treatment | Replicate | | $F_7 = 13.18$, $p = 0.0001$ | lsmeans (Bonferroni correction):<br>GF vs Biotin: $p < 0.0001$<br>GF vs Choline: $p = 1.0$<br>GF vs Folic acid: $p = 1.0$<br>GF vs Nicotinic acid: $p = 1.0$<br>GF vs Pyridoxine: $p = 1.0$<br>GF vs Riboflavin: $p = 0.28$<br>GF vs Thiamine: $p = 1.0$<br>Biotin vs Choline: $p = 0.0001$ | Biotin vs Folic acid: $p < 0.0001$<br>Biotin vs Nicotinic acid: $p < 0.0001$<br>Biotin vs Pyridoxine: $p = 0.0001$<br>Biotin vs Riboflavin: $p = 0.020$<br>Biotin vs Thiamine: $p = 0.0001$<br>Choline vs Folic acid: $p = 1.0$<br>Choline vs Nicotinic acid: $p = 1.0$<br>Choline vs Pyridoxine: $p = 1.0$<br>Choline vs Riboflavin: $p = 0.86$<br>Choline vs Thiamine: $p = 1.0$ | Folic acid vs Nicotinic acid: $p = 1.0$<br>Folic acid vs Pyridoxine: $p = 1.0$<br>Folic acid vs Riboflavin: $p = 1.0$<br>Folic acid vs Thiamine: $p = 1.0$<br>Nicotinic acid vs Pyridoxine: $p = 1.0$<br>Nicotinic acid vs Riboflavin: $p = 0.86$<br>Nicotinic acid vs Thiamine: $p = 1.0$<br>Pyridoxine acid vs Riboflavin: $p = 1.0$<br>Pyridoxine acid vs Thiamine: $p = 1.0$<br>Riboflavin acid vs Thiamine: $p = 1.0$ |
| | % blocked | Treatment | Replicate | | $F_7 = 4.04$, $p = 0.00026$ | lsmeans (Bonferroni correction):<br>GF vs Biotin: $p = 0.096$<br>GF vs Choline: $p = 1.0$<br>GF vs Folic acid: $p = 0.062$<br>GF vs Nicotinic acid: $p = 1.0$<br>GF vs Pyridoxine: $p = 1.0$<br>GF vs Riboflavin: $p = 0.32$<br>GF vs Thiamine: $p = 1.0$<br>Biotin vs Choline: $p = 0.096$ | Biotin vs Folic acid: $p = 1.0$<br>Biotin vs Nicotinic acid: $p = 0.17$<br>Biotin vs Pyridoxine: $p = 0.64$<br>Biotin vs Riboflavin: $p = 1.0$<br>Biotin vs Thiamine: $p = 1.0$<br>Choline vs Folic acid: $p = 0.062$<br>Choline vs Nicotinic acid: $p = 1.0$<br>Choline vs Pyridoxine: $p = 1.0$<br>Choline vs Riboflavin: $p = 0.32$<br>Choline vs Thiamine: $p = 1.0$ | Folic acid vs Nicotinic acid: $p = 0.11$<br>Folic acid vs Pyridoxine: $p = 0.40$<br>Folic acid vs Riboflavin: $p = 1.0$<br>Folic acid vs Thiamine: $p = 0.75$<br>Nicotinic acid vs Pyridoxine: $p = 1.0$<br>Nicotinic acid vs Riboflavin: $p = 0.62$<br>Nicotinic acid vs Thiamine: $p = 1.0$<br>Pyridoxine acid vs Riboflavin: $p = 1.0$<br>Pyridoxine acid vs Thiamine: $p = 1.0$<br>Riboflavin acid vs Thiamine: $p = 1.0$ |

| Analysis | Response variable | Predictor | Random effect | Sample Size | Test result | Comparisons |
|---|---|---|---|---|---|---|
| **Figure 3B** | | | | | | |
| Glmm (binomial) | Sex | Treatment | Replicate | **Replicate A**: GF (n=24); Folic acid (n=24); Choline (n=24); Thiamine (n=24); Nicotinic acid (n=24); Biotin (n=24); Pyridoxine (n=24) **Replicate B**: GF (n=24); Folic acid (n=24); Choline (n=24); Thiamine (n=24); Nicotinic acid (n=24); Biotin (n=24); Riboflavin (n=24); Pyridoxine (n=24) **Replicate C**: GF (n=24); Folic acid (n=24); Choline (n=24); Thiamine (n=24); Nicotinic acid (n=24); Biotin (n=24); Riboflavin (n=24); Pyridoxine (n=24) | $F_7 = 0.63$, $p = 0.73$ | |
| **Figure 3C** | | | | | | |
| Glmm (binomial) | % adults | Treatment | Replicate | **Replicate A**: GF males (n=190); GF females (n=85); biotin 1x males (n=128); biotin 1x females (n=87); biotin 4x males (n=127); biotin 4x females (n=86); biotin 8x males (n=128); biotin 8x females (n=92); biotin 20x males (n=126); biotin 20x females (n=90) **Replicate B**: GF males (n=60); GF females (n=16); GF+folic acid males (n=59); GF+folic acid females (n=16); biotin 1x males (n=60); biotin 1x females (n=24); biotin 4x males (n=57); biotin 4x females (n=23); biotin 8x males (n=57); biotin 8x females (n=24); biotin 20x males (n=59); biotin 20x females (n=24) **Replicate C**: GF males (n=100); GF females (n=93); GF+folic acid males (n=100); GF+folic acid females (n=65); biotin 1x males (n=100); biotin 1x females (n=81); biotin 4x males (n=98); biotin 4x females (n=79); biotin 8x males (n=96); biotin 8x females (n=84); biotin 20x males (n=110); biotin 20x females (n=78) **Replicate D**: GF males (n=112); GF females (n=45); GF+folic acid males (n=149); GF+folic acid females (n=72); biotin 1x males (n=173); biotin 1x females (n=70); biotin 4x males (n=102); biotin 4x females (n=73); biotin 8x males (n=128); biotin 8x females (n=75); biotin 20x males (n=126); biotin 20x females (n=69) | $F_5 = 38.34$, $p < 0.0001$ | lsmeans (Bonferroni correction):<br>GF vs GF+folic acid: $p = 1.0$<br>GF vs Biotinx1: $p < 0.001$<br>GF vs Biotinx4: $p = 0.039$<br>GF vs Biotinx8: $p = 0.025$<br>GF vs Biotinx20: $p < 0.0001$<br>GF+folic acid vs Biotinx1: $p < 0.0001$<br>GF+folic acid vs Biotinx4: $p = 0.54$<br>GF+folic acid vs Biotinx8: $p = 0.012$<br>GF+folic acid vs Biotinx20: $p < 0.0001$<br>Biotinx1 vs Biotinx4: $p = 0.027$<br>Biotinx1 vs Biotinx8: $p < 0.0001$<br>Biotinx1 vs Biotinx20: $p < 0.0001$<br>Biotinx4 vs Biotinx8: $p < 0.0001$<br>Biotinx4 vs Biotinx20: $p < 0.0001$<br>Biotinx8 vs Biotinx20: $p < 0.0001$ |
| | % dead | Treatment | Replicate | | $F_5 = 26.74$, $p < 0.0001$ | lsmeans (Bonferroni correction):<br>GF vs GF+folic acid: $p = 1.0$<br>GF vs Biotinx1: $p < 0.001$<br>GF vs Biotinx4: $p = 0.72$<br>GF vs Biotinx8: $p = 0.10$<br>GF vs Biotinx20: $p < 0.0001$<br>GF+folic acid vs Biotinx1: $p = 0.0001$<br>GF+folic acid vs Biotinx4: $p = 1.0$<br>GF+folic acid vs Biotinx8: $p = 0.065$<br>GF+folic acid vs Biotinx20: $p < 0.0001$<br>Biotinx1 vs Biotinx4: $p = 0.0049$<br>Biotinx1 vs Biotinx8: $p < 0.0001$<br>Biotinx1 vs Biotinx20: $p < 0.0001$<br>Biotinx4 vs Biotinx8: $p = 0.0001$<br>Biotinx4 vs Biotinx20: $p < 0.0001$<br>Biotinx8 vs Biotinx20: $p = 0.25$ |
| | % blocked | Treatment | Replicate | | $F_5 = 2.16$, $p = 0.054$ | |

| Analysis | Response variable | Predictor | Random effect | Sample Size | Test result | Comparisons |
|---|---|---|---|---|---|---|
| **Figure 3D** | | | | | | |
| Glmm (binomial) | % adults | Treatment | Replicate | **Replicate A**: GF males (n=180); biotin 1x males (n=128); biotin 4x males (n=127); biotin 8x males (n=128); biotin 20x males (n=126)<br>**Replicate B**: GF males (n=60); GF+folic acid males (n=59); biotin 1x males (n=60); biotin 4x males (n=57); biotin 8x males (n=57); biotin 20x males (n=59)<br>**Replicate C**: GF males (n=100); GF+folic acid males (n=100); biotin 1x males (n=100); biotin 4x males (n=98); biotin 8x males (n=96); biotin 20x males (n=110)<br>**Replicate D**: GF males (n=112); GF+folic acid males (n=149); biotin 1x males (n=173); biotin 4x males (n=102); biotin 8x males (n=128); biotin 20x males (n=126) | $F_5 = 28.33$, $p < 0.0001$ | lsmeans (Bonferroni correction):<br>GF vs GF+folic acid: $p = 1.0$<br>GF vs Biotinx1: $p < 0.001$<br>GF vs Biotinx4: $p = 1.0$<br>GF vs Biotinx8: $p = 0.019$<br>GF vs Biotinx20: $p < 0.0001$<br>GF+folic acid vs Biotinx1: $p < 0.0001$<br>GF+folic acid vs Biotinx4: $p = 1.0$<br>GF+folic acid vs Biotinx8: $p = 0.13$<br>GF+folic acid vs Biotinx20: $p < 0.0001$<br>Biotinx1 vs Biotinx4: $p < 0.0001$<br>Biotinx1 vs Biotinx8: $p < 0.0001$<br>Biotinx1 vs Biotinx20: $p < 0.0001$<br>Biotinx4 vs Biotinx8: $p = 0.072$<br>Biotinx4 vs Biotinx20: $p < 0.0001$<br>Biotinx8 vs Biotinx20: $p = 0.013$ |
| | % dead | Treatment | Replicate | | $F_5 = 18.43$, $p < 0.0001$ | lsmeans (Bonferroni correction):<br>GF vs GF+folic acid: $p = 1.0$<br>GF vs Biotinx1: $p < 0.001$<br>GF vs Biotinx4: $p = 1.0$<br>GF vs Biotinx8: $p = 0.19$<br>GF vs Biotinx20: $p = 0.022$<br>GF+folic acid vs Biotinx1: $p = 0.0001$<br>GF+folic acid vs Biotinx4: $p = 1.0$<br>GF+folic acid vs Biotinx8: $p = 0.59$<br>GF+folic acid vs Biotinx20: $p = 0.017$<br>Biotinx1 vs Biotinx4: $p < 0.0001$<br>Biotinx1 vs Biotinx8: $p < 0.0001$<br>Biotinx1 vs Biotinx20: $p < 0.0001$<br>Biotinx4 vs Biotinx8: $p = 0.63$<br>Biotinx4 vs Biotinx20: $p = 0.014$<br>Biotinx8 vs Biotinx20: $p = 1.0$ |
| | % blocked | Treatment | Replicate | | $F_5 = 2.55$, $p = 0.025$ | lsmeans (Bonferroni correction):<br>GF vs GF+folic acid: $p = 1.0$<br>GF vs Biotinx1: $p = 1.0$<br>GF vs Biotinx4: $p = 1.0$<br>GF vs Biotinx8: $p = 1.0$<br>GF vs Biotinx20: $p = 0.22$<br>GF+folic acid vs Biotinx1: $p = 1.0$<br>GF+folic acid vs Biotinx4: $p = 1.0$<br>GF+folic acid vs Biotinx8: $p = 1.0$<br>GF+folic acid vs Biotinx20: $p = 1.0$<br>Biotinx1 vs Biotinx4: $p = 1.0$<br>Biotinx1 vs Biotinx8: $p = 1.0$<br>Biotinx1 vs Biotinx20: $p = 0.030$<br>Biotinx4 vs Biotinx8: $p = 1.0$<br>Biotinx4 vs Biotinx20: $p = 0.22$<br>Biotinx8 vs Biotinx20: $p = 1.0$ |

| Analysis | Response variable | Predictor | Random effect | Sample Size | Test result | Comparisons |
|---|---|---|---|---|---|---|
| **Figure 3E** | | | | | | |
| Glmm (binomial) | % adults | Treatment | Replicate | **Replicate A**: GF females (n=85); biotin 1x females (n=87); biotin 4x females (n=86); biotin 8x females (n=52); biotin 20x females (n=90)<br>**Replicate B**: GF females (n=16); GF+folic acid females (n=16); biotin 1x females (n=24); biotin 4x females (n=23); biotin 8x females (n=24); biotin 20x females (n=24)<br>**Replicate C**: GF females (n=93); GF+folic acid females (n=65); biotin 1x females (n=81); biotin 4x females (n=79); biotin 8x females (n=84); biotin 20x females (n=78)<br>**Replicate D**: GF females (n=45); GF+folic acid females (n=72); biotin 1x females (n=70); biotin 4x females (n=73); biotin 8x females (n=75); biotin 20x females (n=69) | $F_1 = 16.77$, $p < 0.0001$ | lsmeans (Bonferroni correction):<br>GF vs GF+folic acid: $p = 1.0$<br>GF vs Biotinx1: $p = 0.038$<br>GF vs Biotinx4: $p = 0.0001$<br>GF vs Biotinx8: $p = 1.0$<br>GF vs Biotinx20: $p = 0.0018$<br>GF+folic acid vs Biotinx1: $p = 1.0$<br>GF+folic acid vs Biotinx4: $p = 0.25$<br>GF+folic acid vs Biotinx8: $p = 0.13$<br>GF+folic acid vs Biotinx20: $p = 0.0001$<br>Biotinx1 vs Biotinx4: $p = 1.0$<br>Biotinx1 vs Biotinx8: $p = 0.0004$<br>Biotinx1 vs Biotinx20: $p < 0.0001$<br>Biotinx4 vs Biotinx8: $p = 0.0001$<br>Biotinx4 vs Biotinx20: $p < 0.0001$<br>Biotinx8 vs Biotinx20: $p = 0.049$ |
| | % dead | Treatment | Replicate | | $F_1 = 12.45$, $p = 0.0003$ | lsmeans (Bonferroni correction):<br>GF vs GF+folic acid: $p = 1.0$<br>GF vs Biotinx1: $p = 0.068$<br>GF vs Biotinx4: $p = 0.034$<br>GF vs Biotinx8: $p = 1.0$<br>GF vs Biotinx20: $p = 0.013$<br>GF+folic acid vs Biotinx1: $p = 1.0$<br>GF+folic acid vs Biotinx4: $p = 1.0$<br>GF+folic acid vs Biotinx8: $p = 0.12$<br>GF+folic acid vs Biotinx20: $p = 0.0009$<br>Biotinx1 vs Biotinx4: $p = 1.0$<br>Biotinx1 vs Biotinx8: $p = 0.0001$<br>Biotinx1 vs Biotinx20: $p = 0.0001$<br>Biotinx4 vs Biotinx8: $p < 0.0001$<br>Biotinx4 vs Biotinx20: $p = 0.0001$<br>Biotinx8 vs Biotinx20: $p = 0.76$ |
| | % blocked | Treatment | Replicate | | $F_1 = 1.28$, $p = 0.27$ | |
| **Figure 4A** | | | | | | |
| Glmm (Type III Analysis of Variance Table with Satterthwaite's method) | mean difference | Treatment | / | **Replicate A**: Treatment 1x, day 0 (n=6), day 1 (n=6), day 3 (n=6), day 6 (n=6), day 7 (n=6), day 8 (n=6), day 9 (n=6), day 10 (n=6). Treatment 0.01x, day 0 (n=6), day 1 (n=6), day 2 (n=6), day 3 (n=6), day 6, (n=6), day 7 (n=6), day 8 (n=6), day 9 (n=6), day 10 (n=6).<br>**Replicate B**: Treatment 1x, day 0 (n=6), day 1 (n=6), day 2 (n=6), day 3 (n=6), day 6, (n=6), day 7 (n=6), day 8 (n=6), day 9 (n=6). Treatment 0.01x, day 0 (n=6), day 1 (n=6), day 2 (n=6), day 3 (n=6), day 6, (n=6), day 7 (n=6), day 8 (n=6), day 9 (n=6).<br>**Replicate C**: Treatment 1x, day 0 (n=6), day 1 (n=6), day 2 (n=6), day 3 (n=6), day 6, (n=6), day 7 (n=6), day 8 (n=6), day 9 (n=6). Treatment 0.01x, day 0 (n=6), day 1 (n=6), day 2 (n=6), day 3 (n=6), day 6, (n=6), day 7 (n=6), day 8 (n=6), day 9 (n=6).<br>**Replicate D**: Treatment 1x, day 0 (n=6), day 1 (n=6), day 2 (n=6), day 3 (n=6), day 6, (n=6), day 7 (n=6), day 10 (n=6). Treatment 0.01x, day 0 (n=6), day 1 (n=6), day 2 (n=6), day 3 (n=6), day 6, (n=6), day 7 (n=6), day 10 (n=6).<br>**Replicate E**: Treatment 1x, day 0 (n=6), day 1 (n=6), day 2 (n=6), day 3 (n=6), day 6, (n=6), day 7 (n=6), day 10 (n=6). Treatment 0.01x, day 0 (n=6), day 1 (n=6), day 2 (n=6), day 3 (n=6), day 6, (n=6), day 7 (n=6), day 10 (n=6). | 1x vs 0.01 x, day 0<br>F = 1.1718, p = 0.28387<br>1x vs 0.01 x, day 1<br>F = 44.806, $p < 0.0001$<br>1x vs 0.01 x, day 2<br>F = 20.775, $p < 0.0001$<br>1x vs 0.01 x, day 3<br>F = 30.31, $p < 0.0001$<br>1x vs 0.01 x, day 6<br>F = 54.307, $p = 0.0001$<br>1x vs 0.01 x, day 7<br>F = 36.18, $p = 0.0001$<br>1x vs 0.01 x, day 8<br>F = 6.2088, $p = 0.0127$<br>1x vs 0.01 x, day 9<br>F = 10.235, $p = 0.00138$<br>1x vs 0.01 x, day 10<br>F = 27.115, $p < 0.0001$ | |
| **Figure 4B** | | | | | | |
| Glmm (binomial) | % adults | Treatment | Replicate | **Replicate A**: Treatment 1x, larvae (n=24). Treatment 0.01x, larvae (n=24)<br>**Replicate B**: Treatment 1x, larvae (n=24). Treatment 0.01x, larvae (n=24)<br>**Replicate C**: Treatment 1x, larvae (n=24). Treatment 0.01x, larvae (n=24)<br>**Replicate D**: Treatment 1x, larvae (n=48). Treatment 0.01x, larvae (n=48)<br>**Replicate E**: Treatment 1x, larvae (n=48). Treatment 0.01x, larvae | F = 76.218, $p < 0.0001$ | lsmeans (Bonferroni correction)<br>1x vs 0.01x, $p < 0.0001$ |
| | % dead | Treatment | Replicate | | F = 54.708, $p = 0.0001$ | lsmeans (Bonferroni correction)<br>1x vs 0.01x, $p < 0.0001$ |
| | % blocked | Treatment | Replicate | | F = 4.3810, $p = 0.000131$ | lsmeans (Bonferroni correction)<br>1x vs 0.01x, $p = 0.0382$ |

| Analysis | Response variable | Predictor | Random effect | Sample Size | Test result | Comparisons |
|---|---|---|---|---|---|---|
| **Figure 4C** | | | | | | |
| Glmm (binomial) | Sex | Treatment | Replicate | **Replicate A:** Treatment 1x, larvae (n=24). Treatment 0.01x, larvae (n=24) **Replicate B:** Treatment 1x, larvae (n=24). Treatment 0.01x, larvae | F = 3.8, *p* = 0.043 | |
| **Figure 4D** | | | | | | |
| Glmm (Type III Analysis of Variance Table with Satterthwaite's method) | Time | Treatment | Replicate | **Replicate A:** Treatment 1x, larvae (n=24). Treatment 0.01x, larvae (n=24) **Replicate B:** Treatment 1x, larvae (n=24). Treatment 0.01x, larvae (n=24) **Replicate C:** Treatment 1x, larvae (n=24). Treatment 0.01x, larvae (n=24) **Replicate D:** Treatment 1x, larvae (n=48). Treatment 0.01x, larvae (n=48) **Replicate E:** Treatment 1x, larvae (n=48). Treatment 0.01x, larvae (n=48) | F = 45.292, *p* = 0.0115 | |
| **Figure 4E** | | | | | | |
| Glmm (binomial) | % adults | Treatment | Replicate | **Replicate A:** wt strain, larvae (n=120). ΔscUA strain, larvae (n=120). ΔmmmA strain, larvae (n=120). **Replicate B:** wt strain, larvae (n=120). ΔscUA strain, larvae (n=120). ΔmmmA strain, larvae (n=120). **Replicate C:** wt strain, larvae (n=120). ΔscUA strain, larvae (n=120). ΔmmmA strain, larvae (n=120). **Replicate D:** wt strain, larvae (n=120). ΔscUA strain, larvae (n=120). ΔmmmA strain, larvae (n=120). **Replicate E:** wt strain, larvae (n=120). ΔscUA strain, larvae (n=120). ΔmmmA strain, larvae (n=120). | F = 4.9097<br>wt vs ΔscUA: *p* = 0.0019<br>wt vs ΔmmmA: *p* = 0.14743 | |
| | % dead | Treatment | Replicate | | F = 3.4591<br>wt vs ΔscUA: *p* = 0.00973<br>wt vs ΔmmmA: *p* = 0.11306 | |
| | % blocked | Treatment | Replicate | | F = 1.2511<br>wt vs ΔscUA: *p* = 0.134<br>wt vs ΔmmmA: *p* = 0.583 | |
| **Figure 4F** | | | | | | |
| Glmm (binomial) | Sex | Treatment | Replicate | **Replicate A:** wt strain, larvae (n=120). ΔscUA strain, larvae (n=120). ΔmmmA strain, larvae (n=120). **Replicate B:** wt strain, larvae (n=120). ΔscUA strain, larvae (n=120). ΔmmmA strain, larvae (n=120). **Replicate C:** wt strain, larvae (n=120). ΔscUA strain, larvae (n=120). ΔmmmA strain, larvae (n=120). **Replicate D:** wt strain, larvae (n=120). ΔscUA strain, larvae (n=120). ΔmmmA strain, larvae (n=120). **Replicate E:** wt strain, larvae (n=120). ΔscUA strain, larvae (n=120). ΔmmmA strain, larvae (n=120). | F = 3.2628, *p* = 0.03714<br>wt vs ΔscUA: *p* = 0.0847<br>wt vs ΔmmmA: *p* = 0.0147 | |
| **Figure S1** | | | | | | |
| Glmm (binomial) | % adults | Treatment | Replicate | **Replicate A:** GF (n=48); VIT1x (n=48) **Replicate B:** GF (n=48); VIT1x (n=48) **Replicate C:** GF (n=48); VIT1x (n=48) | $F_1$ = 0.35, *p* = 0.55 | |
| | % dead | Treatment | Replicate | | $F_1$ = 0.11, *p* = 0.74 | |
| | % blocked | Treatment | Replicate | | $F_1$ = 0, *p* = 1.0 | |
| **Figure S4A** | | | | | | |
| Log-rank (Mantel-Cox) test | % pupae GF | Sex | / | **Replicate A:** GF males (n=150); GF females (n=85); biotin 1x males (n=128); biotin 1x females (n=87); biotin 4x males (n=127); biotin 4x females (n=86); biotin 8x males (n=128); biotin 8x females (n=92); biotin 20x males (n=128); biotin 20x females (n=90) **Replicate B:** GF males (n=60); GF females (n=36); GF+folic acid males (n=59); GF+folic acid females (n=16); biotin 1x males (n=60); biotin 1x females (n=24); biotin 4x males (n=57); biotin 4x females (n=33); biotin 8x males (n=57); biotin 8x females (n=24); biotin 20x males (n=58); biotin 20x females (n=24) **Replicate C:** GF males (n=100); GF females (n=69); GF+folic acid males (n=100); GF+folic acid females (n=65); biotin 1x males (n=100); biotin 1x females (n=81); biotin 4x males (n=98); biotin 4x females (n=73); biotin 8x males (n=96); biotin 8x females (n=84); biotin 20x males (n=110); biotin 20x females (n=78) **Replicate D:** GF males (n=112); GF females (n=45); GF+folic acid males (n=149); GF+folic acid females (n=72); biotin 1x males (n=173); biotin 1x females (n=70); biotin 4x males (n=102); biotin 4x females (n=73); biotin 8x males (n=128); biotin 8x females (n=75); biotin 20x males (n=126); biotin 20x females (n=65) | $\chi_1$ = 26.83, *p* < 0.0001 | / |
| | % pupae GF+folic acid | Sex | / | | $\chi_1$ = 0.53, *p* = 0.46 | / |
| | % pupae biotin 1x | Sex | / | | $\chi_1$ = 21.68, *p* = 0.0001 | / |
| | % pupae biotin 4x | Sex | / | | $\chi_1$ = 0.0030, *p* = 0.96 | / |
| | % pupae biotin 8x | Sex | / | | $\chi_1$ = 5.44, *p* = 0.020 | / |
| | % pupae biotin 20x | Sex | / | | $\chi_1$ = 0.69, *p* = 0.57 | / |

| Analysis | Response variable | Predictor | Random effect | Sample Size | Test result | Comparisons |
|---|---|---|---|---|---|---|
| **Figure S4B** | | | | | | |
| Log-rank (Mantel-Cox) test | % adults GF | Sex | / | **Replicate A:** GF males (n=100); GF females (n=85); biotin 1x males (n=128); biotin 1x females (n=87); biotin 4x males (n=127); biotin 4x females (n=88); biotin 8x males (n=128); biotin 8x females (n=92); biotin 20x males (n=126); biotin 20x females (n=90) **Replicate B:** GF males (n=80); GF females (n=18); GF+folic acid males (n=59); GF+folic acid females (n=36); biotin 1x males (n=60); biotin 1x females (n=24); biotin 4x males (n=57); biotin 4x females (n=23); biotin 8x males (n=57); biotin 8x females (n=24); biotin 20x males (n=59); biotin 20x females (n=24) **Replicate C:** GF males (n=100); GF females (n=93); GF+folic acid males (n=100); GF+folic acid females (n=65); biotin 1x males (n=100); biotin 1x females (n=81); biotin 4x males (n=88); biotin 4x females (n=78); biotin 8x males (n=98); biotin 8x females (n=84); biotin 20x males (n=110); biotin 20x females (n=78) **Replicate D:** GF males (n=132); GF females (n=45); GF+folic acid males (n=148); GF+folic acid females (n=72); biotin 1x males (n=173); biotin 1x females (n=70); biotin 4x males (n=102); biotin 4x females (n=73); biotin 8x males (n=128); biotin 8x females (n=75); biotin 20x males (n=126); biotin 20x females (n=68) | $\chi^2_1 = 0.045$, $p = 0.83$ | / |
| | % adults GF+folic acid | Sex | / | | $\chi^2_1 = 12.08$, $p = 0.0005$ | / |
| | % adults biotin 1x | Sex | / | | $\chi^2_1 = 1.18$, $p = 0.28$ | / |
| | % adults biotin 4x | Sex | / | | $\chi^2_1 = 41.17$, $p < 0.0001$ | / |
| | % adults biotin 8x | Sex | / | | $\chi^2_1 = 6.02$, $p = 0.014$ | / |
| | % adults biotin 20x | Sex | / | | $\chi^2_1 = 5.12$, $p = 0.024$ | / |
| **Figure S5A** | | | | | | |
| Glmm (Type III Analysis of Variance Table with Satterthwaite's method) | Time | Treatment | Replicate | Replicate A: Treatment 1x, day 0 (n=3), day 1 (n=3), day 3 (n=3), day 6 (n=3), day 7 (n=3), day 8 (n=3), day 9 (n=3), day 10 (n=3). Treatment 0.1x, day 0 (n=3), day 1 (n=3), day 2 (n=3), day 3 (n=3), day 6, (n=3), day 7 (n=3), day 8 (n=3), day 9 (n=3), day 10 (n=3) Replicate B: Treatment 1x, day 0 (n=3), day 1 (n=3), day 2 (n=3), day 3 (n=3), day 6, (n=3), day 7 (n=3), day 8 (n=3), day 9 (n=3). Treatment 0.1x, day 0 (n=3), day 1 (n=3), day 2 (n=3), day 3 (n=3), day 6, (n=3), day 7 (n=3), day 8 (n=3), day 9 (n=3) Replicate C: Treatment 1x, day 0 (n=3), day 1 (n=3), day 1 (n=3), day 3 (n=3), day 6, (n=3), day 7 (n=3), day 8 (n=3), day 9 (n=3). Treatment 0.1x, day 0 (n=3), day 1 (n=3), day 2 (n=3), day 3 (n=3), day 6, (n=3), day 7 (n=3), day 8 (n=3), day 9 (n=3). | 1x vs 0.1 x, day 0<br>$F = 0.4488$, $p = 0.5125$ | |
| | | | | | 1x vs 0.1 x, day 1<br>$F = 26.162$, $p < 0.0001$ | |
| | | | | | 1x vs 0.1 x, day 2<br>$F = 16.807$, $p < 0.0001$ | |
| | | | | | 1x vs 0.1 x, day 3<br>$F = 9.6785$, $p = 0.001864$ | |
| | | | | | 1x vs 0.1 x, day 6<br>$F = 9.0583$, $p = 0.002618$ | |
| | | | | | 1x vs 0.1 x, day 7<br>$F = 0.205$, $p = 0.6507$ | |
| | | | | | 1x vs 0.1 x, day 8<br>$F = 0.0959$, $p = 0.7568$ | |
| | | | | | 1x vs 0.1 x, day 9<br>$F = 4.5143$, $p = 0.03361$ | |
| | | | | | 1x vs 0.1 x, day 10<br>n/a | |
| **Figure S5B** | | | | | | |
| Glmm (binomial) | % adults | Treatment | Replicate | **Replicate A:** Treatment 1x, larvae (n=24). Treatment 0.1x, larvae (n=24) **Replicate B:** Treatment 1x, larvae (n=24). Treatment 0.1x, larvae (n=24) **Replicate C:** Treatment 1x, larvae (n=24). Treatment 0.1x, larvae (n=24) | $F = 7.2695$, $p = 0.006121$ | |
| | % dead | Treatment | Replicate | | $F = 1.2501$, $p = 0.262$ | |
| | % bloated | Treatment | Replicate | | $F = 4.648$, $p = 0.03105$ | |
| **Figure S5C** | | | | | | |
| Glmm (binomial) | Sex | Treatment | Replicate | **Replicate A:** Treatment 1x, larvae (n=24). Treatment 0.1x, larvae (n=24) **Replicate B:** Treatment 1x, larvae (n=24). Treatment 0.1x, larvae (n=24) **Replicate C:** Treatment 1x, larvae (n=24). Treatment 0.1x, larvae (n=24) | $F = 0.9739$, $p = 0.3237$ | |
| **Figure S5D** | | | | | | |
| Glmm (Type III Analysis of Variance Table with Satterthwaite's method) | Time | Treatment | Replicate | **Replicate A:** Treatment 1x, larvae (n=24). Treatment 0.1x, larvae (n=24) **Replicate B:** Treatment 1x, larvae (n=24). Treatment 0.1x, larvae (n=24) **Replicate C:** Treatment 1x, larvae (n=24). Treatment 0.1x, larvae (n=24) | $F = 6.5262$, $p = 0.0115$ | |

**Supplementary Table S2. List of genes differentially expressed between males and females in germ-free conditions.** For each gene, Vectorbase ID, expression value in males compared to females (log2FoldChange), *p*adj, Vectorbase description, and transcript type are indicated. The last column indicates whether the same gene was identified as over- or under-expressed in male larvae in (Matthews et al. 2018).

| ID | log2FoldChange | *padj* | Transcript product | Transcript type | Present in Matthews et al with the same sign |
|---|---|---|---|---|---|
| AAEL003771 | -8.295687137 | 0.019909552 | trithorax protein ash2 | protein_coding_gene | |
| AAEL024436 | -7.816682861 | 0.046563698 | unspecified product | protein_coding_gene | |
| AAEL024463 | -3.069825309 | 0.048126638 | unspecified product | protein_coding_gene | |
| **AAEL001822** | **-2.915032564** | **2.38672E-13** | **glucosyl/glucuronosyl transferases** | **protein_coding_gene** | Yes |
| **AAEL004805** | **-2.670630469** | **1.59585E-25** | **potassium-dependent sodium-calcium exchanger, putative** | **protein_coding_gene** | Yes |
| **AAEL010789** | **-2.437277909** | **1.44998E-27** | **unspecified product** | **protein_coding_gene** | Yes |
| AAEL007305 | -2.07283861 | 0.013687482 | unspecified product | protein_coding_gene | |
| AAEL010062 | -1.736864882 | 0.00127328 | unspecified product | protein_coding_gene | |
| AAEL017646 | -1.626913315 | 1.08852E-08 | U1 spliceosomal RNA | ncRNA_gene | |
| AAEL017799 | -1.549481624 | 3.92516E-05 | U1 spliceosomal RNA | ncRNA_gene | |
| AAEL017691 | -1.518186025 | 2.83258E-06 | U1 spliceosomal RNA | ncRNA_gene | |
| AAEL025856 | -1.480165441 | 0.000375634 | unspecified product | protein_coding_gene | |
| AAEL021440 | -1.396719083 | 0.000744631 | unspecified product | ncRNA_gene | |
| AAEL009601 | -1.381227273 | 6.17919E-05 | pyridoxine kinase | protein_coding_gene | Yes |
| AAEL014357 | -1.375674629 | 0.018958086 | unspecified product | protein_coding_gene | |
| AAEL020258 | -1.205017614 | 8.93111E-05 | unspecified product | protein_coding_gene | |
| AAEL005821 | -1.047163745 | 0.001440823 | alanyl aminopeptidase | protein_coding_gene | Yes |
| AAEL017826 | -0.988811346 | 8.45295E-07 | U1 spliceosomal RNA | ncRNA_gene | |
| AAEL004188 | -0.96318036 | 0.017351326 | unspecified product | protein_coding_gene | Yes |
| AAEL028652 | -0.9365906 | 3.92516E-05 | unspecified product | ncRNA_gene | |
| AAEL023559 | -0.929041606 | 0.002171398 | unspecified product | protein_coding_gene | Yes |
| AAEL028907 | -0.846734961 | 0.00556439 | unspecified product | ncRNA_gene | |
| AAEL005616 | -0.844406292 | 0.028419036 | trypsin | protein_coding_gene | |
| AAEL009537 | -0.84378579 | 0.002160461 | unspecified product | protein_coding_gene | |
| AAEL008380 | -0.841141496 | 0.002478452 | unspecified product | protein_coding_gene | Yes |
| AAEL005702 | -0.733722783 | 0.005111812 | unspecified product | protein_coding_gene | |
| AAEL000931 | -0.691037776 | 0.032076329 | Alkaline phosphatase | protein_coding_gene | Yes |
| AAEL009150 | 0.621817125 | 0.03820079 | unspecified product | protein_coding_gene | Yes |
| AAEL008451 | 0.782356219 | 0.03866026 | alpha-amylase | protein_coding_gene | |
| AAEL008789 | 0.804032867 | 0.033012268 | apolipophorin-III, putative | protein_coding_gene | |
| AAEL004978 | 0.806425849 | 0.00767461 | DEAD box ATP-dependent RNA helicase | protein_coding_gene | Yes |
| AAEL011031 | 0.849670541 | 0.008544662 | unspecified product | protein_coding_gene | Yes |
| AAEL011889 | 0.852421565 | 0.01855757 | serine-type enodpeptidase, | protein_coding_gene | |
| AAEL008708 | 0.870414126 | 0.002122255 | lysosomal pro-X carboxypeptidase, putative | protein_coding_gene | |
| AAEL013484 | 0.879055901 | 0.015788794 | unspecified product | protein_coding_gene | Yes |
| AAEL015247 | 0.889526773 | 0.003811927 | unspecified product | protein_coding_gene | |
| AAEL027166 | 0.898050756 | 0.006611833 | unspecified product | protein_coding_gene | Yes |
| AAEL012558 | 0.909596157 | 0.046563698 | serine collagenase 1 precursor, putative | protein_coding_gene | |
| AAEL021888 | 1.013941359 | 0.001021322 | unspecified product | protein_coding_gene | |
| AAEL008207 | 1.022171954 | 0.024935899 | serine protease | protein_coding_gene | Yes |
| AAEL019919 | 1.028249018 | 0.004463092 | unspecified product | protein_coding_gene | |
| AAEL020614 | 1.031255967 | 0.000369921 | unspecified product | protein_coding_gene | Yes |
| AAEL003905 | 1.037437949 | 0.008078978 | alkaline phosphatase | protein_coding_gene | |
| AAEL005609 | 1.057387147 | 0.017545294 | trypsin | protein_coding_gene | |
| AAEL004344 | 1.168985667 | 0.002478452 | zinc finger protein | protein_coding_gene | |
| AAEL006627 | 1.170903873 | 1.5634E-05 | serine-type enodpeptidase, | protein_coding_gene | Yes |
| AAEL008598 | 1.215471286 | 0.000399915 | unspecified product | protein_coding_gene | |
| AAEL017056 | 1.266710284 | 0.000164633 | Peptidoglycan Recognition Protein (Short) | protein_coding_gene | Yes |
| AAEL001897 | 1.379110817 | 0.01536703 | unspecified product | protein_coding_gene | |
| AAEL005431 | 1.474760847 | 0.013687482 | Clip-Domain Serine Protease family B. | protein_coding_gene | |
| AAEL002670 | 1.534857547 | 0.016213222 | AMP dependent ligase | protein_coding_gene | |
| AAEL002680 | 1.559770116 | 0.046563698 | AMP dependent ligase | protein_coding_gene | |
| AAEL007039 | 1.598761821 | 1.86007E-08 | peptidoglycan recognition protein (short) | protein_coding_gene | Yes |
| AAEL008604 | 1.604397372 | 0.003683224 | zinc carboxypeptidase | protein_coding_gene | Yes |
| AAEL011169 | 2.110355525 | 3.25274E-19 | hexamerin 2 beta | protein_coding_gene | |
| AAEL012018 | 2.224734632 | 0.010496751 | unspecified product | protein_coding_gene | |
| AAEL019574 | 2.285204913 | 7.81614E-05 | unspecified product | protein_coding_gene | |
| AAEL013757 | 2.409701453 | 5.34455E-20 | hexamerin 2 beta | protein_coding_gene | |
| AAEL013983 | 2.646392815 | 2.64812E-09 | hexamerin 2 beta | protein_coding_gene | Yes |
| AAEL023288 | 2.762499229 | 0.021713334 | unspecified product | protein_coding_gene | |
| **AAEL005606** | **3.048159131** | **0.029159311** | **unspecified product** | **protein_coding_gene** | |
| AAEL013981 | 3.610094666 | 0.01784989 | hexamerin 2 beta | protein_coding_gene | Yes |
| **AAEL013692** | **3.927779529** | **1.14294E-12** | **PIWI** | **protein_coding_gene** | Yes |
| AAEL004316 | 4.907301955 | 0.000742369 | unspecified product | protein_coding_gene | |
| **AAEL022912** | **5.533191955** | **1.28854E-14** | **Male determiner protein Nix** | **protein_coding_gene** | Yes |
| AAEL011582 | 5.567452042 | 3.5922E-06 | unspecified product | protein_coding_gene | |
| AAEL010008 | 5.92893559 | 0.01536703 | crotonobetainyl-CoA-hydratase, putative | protein_coding_gene | |
| **AAEL022112** | **7.211578145** | **5.4287E-06** | **unspecified product** | **ncRNA_gene** | |
| **AAEL022711** | **10.10632484** | **5.88711E-18** | **unspecified product** | **ncRNA_gene** | Yes |

**Supplementary Table S3. List of genes differentially expressed between males and females in vitx8 conditions.** For each gene, Vectorbase ID, expression value in males compared to females (log2FoldChange), *p*adj, Vectorbase description, and transcript type are indicated. The last column indicates whether the same gene was identified as over- or under-expressed in male larvae in (Matthews et al. 2018).

| ID | log2FoldChange | *padj* | Transcript product | Transcript type | Present in Matthews and with the same sign |
|---|---|---|---|---|---|
| AAEL011777 | -5.560831347 | 0.021495448 | Serine Protease Inhibitor (serpin) likely cleavage at K/A. | protein_coding_gene | Yes |
| **AAEL004805** | **-2.648985682** | **2.05779E-07** | **potassium-dependent sodium-calcium exchanger, putative** | **protein_coding_gene** | Yes |
| AAEL028730 | -2.531257596 | 0.000400978 | unspecified product | ncRNA_gene | No |
| **AAEL010789** | **-2.383150143** | **4.8334E-10** | **unspecified product** | **protein_coding_gene** | Yes |
| **AAEL001822** | **-2.27839688** | **0.000200225** | **glucosyl/glucuronosyl transferases** | **protein_coding_gene** | Yes |
| AAEL004564 | 1.948725295 | 0.037936324 | unspecified product | protein_coding_gene | |
| AAEL018263 | 2.302792523 | 0.030091688 | unspecified product | protein_coding_gene | |
| **AAEL013692** | **2.360816866** | **0.002542467** | **PIWI** | **protein_coding_gene** | Yes |
| AAEL008595 | 2.616380998 | 0.037936324 | Protein maelstrom homolog | protein_coding_gene | |
| AAEL010097 | 2.779513254 | 0.016056517 | unspecified product | protein_coding_gene | |
| AAEL001237 | 2.988698973 | 0.037936324 | unspecified product | protein_coding_gene | |
| **AAEL005606** | **3.297367166** | **0.018085638** | **unspecified product** | **protein_coding_gene** | |
| **AAEL022112** | **3.439996092** | **0.013266686** | **unspecified product** | **ncRNA_gene** | |
| AAEL020037 | 3.520829382 | 0.030623181 | unspecified product | protein_coding_gene | |
| **AAEL022912** | **3.749367667** | **1.17663E-07** | **Male determiner protein Nix** | **protein_coding_gene** | Yes |
| **AAEL022711** | **3.85767514** | **8.99411E-19** | **unspecified product** | **ncRNA_gene** | Yes |
| AAEL021838 | 4.28749305 | 1.27962E-06 | Myosin heavy chain | protein_coding_gene | Yes |
| AAEL025512 | 4.513381336 | 0.021495448 | unspecified product | pseudogene | |
| AAEL026034 | 5.003862834 | 0.020028118 | unspecified product | ncRNA_gene | |
| AAEL027534 | 6.945152012 | 0.037936324 | unspecified product | pseudogene | |

**Supplementary Table S4. List of primers used in this study.**

| Primer name | Vectorbase accession | Sequence (5'-3') | Amplicon size | Reference |
|---|---|---|---|---|
| Nix-FOR | AAEL0922912 | GATTTTTGTTTTTGTCGTGCAA | 350 bp | (Hall et al. 2015) |
| Nix-FOR | | AATGCAAGTATCATAGGTCAGCA | | |
| Act-FOR | AAEL011197 | CGTTCGTGACATCAAGGAAA | 175 bp | (Dzaki et al. 2017) |
| Act-REV | | GAACGATGGCTGGAAGAGAG | | |

### Supplemental References

Aryan A, Anderson MAE, Biedler JK, Qi Y, Overcash JM, Naumenko AN, Sharakhova M V., Mao C, Adelman ZN, Tu Z. 2020. Nix alone is sufficient to convert female Aedes aegypti into fertile males and myo-sex is needed for male flight. *Proc Natl Acad Sci U S A* **117**: 17702–17709. /pmc/articles/PMC7395513/ (Accessed September 4, 2023).

Bolger AM, Lohse M, Usadel B. 2014. Trimmomatic: a flexible trimmer for Illumina sequence data. *Bioinformatics* **30**: 2114. /pmc/articles/PMC4103590/ (Accessed September 4, 2023).

Dzaki N, Ramli KN, Azlan A, Ishak IH, Azzam G. 2017. Evaluation of reference genes at different developmental stages for quantitative real-time PCR in Aedes aegypti. *Scientific Reports 2017 7:1* **7**: 1–13. https://www.nature.com/articles/srep43618 (Accessed September 4, 2023).

Hall AB, Basu S, Jiang X, Qi Y, Timoshevskiy VA, Biedler JK, Sharakhova M V., Elahi R, Anderson MAE, Chen XG, et al. 2015. A male-determining factor in the mosquito Aedes aegypti. *Science* **348**: 1268. /pmc/articles/PMC5026532/ (Accessed September 4, 2023).

Kim D, Paggi JM, Park C, Bennett C, Salzberg SL. 2019. Graph-based genome alignment and genotyping with HISAT2 and HISAT-genotype. *Nature Biotechnology 2019 37:8* **37**: 907–915. https://www.nature.com/articles/s41587-019-0201-4 (Accessed September 4, 2023).

Liao Y, Smyth GK, Shi W. 2014. featureCounts: an efficient general purpose program for assigning sequence reads to genomic features. *Bioinformatics* **30**: 923–930. https://dx.doi.org/10.1093/bioinformatics/btt656 (Accessed September 4, 2023).

Love MI, Huber W, Anders S. 2014. Moderated estimation of fold change and dispersion for RNA-seq data with DESeq2. *Genome Biol* **15**: 550. /pmc/articles/PMC4302049/ (Accessed September 4, 2023).

Matthews BJ, Dudchenko O, Kingan SB, Koren S, Antoshechkin I, Crawford JE, Glassford WJ, Herre M, Redmond SN, Rose NH, et al. 2018. Improved reference genome of Aedes aegypti informs arbovirus vector control. *Nature* **563**: 501–507. https://pubmed.ncbi.nlm.nih.gov/30429615/ (Accessed May 17, 2023).

Ram KR, Wolfner MF. 2007. Seminal influences: Drosophila Acps and the molecular interplay between males and females during reproduction. *Integr Comp Biol* **47**: 427–445. https://dx.doi.org/10.1093/icb/icm046 (Accessed September 4, 2023).

Babraham Bioinformatics - FastQC A Quality Control tool for High Throughput Sequence Data. https://www.bioinformatics.babraham.ac.uk/projects/fastqc/ (Accessed September 4, 2023).
